# Generalizable direct protein sequencing with InstaNexus

**DOI:** 10.1101/2025.07.25.666861

**Authors:** Marco Reverenna, Maike Wennekers Nielsen, Darian Stephan Wolff, Elpida Lytra, Pasquale D. Colaianni, Anne Ljungars, Andreas H. Laustsen, Erwin M. Schoof, Jeroen Van Goey, Timothy P. Jenkins, Marie V. Lukassen, Alberto Santos, Konstantinos Kalogeropoulos

## Abstract

Protein-based therapeutics, such as antibodies and nanobodies, are not encoded in reference genomes, challenging their accurate characterization via standard proteomics. Current methods rely on indirect inference, fragmented outputs, and labor-intensive workflows, which hinder functional insights and routine application. Here, we present a generalizable, end-to-end workflow for direct protein sequencing, combining streamlined sample preparation, AI-driven *de novo* peptide sequencing, and tailored assembly to reconstruct contiguous protein sequences. A novel composite scoring framework prioritises longer assemblies and coverage, enhancing accuracy and reducing ambiguity. Validation across diverse protein modalities demonstrates its utility and ability to robustly sequence functionally critical regions of selected proteins. This workflow represents an advance in precision proteomics with promising applications in therapeutic discovery, immune profiling, and protein science.

## Introduction

Accurate protein sequencing remains a fundamental challenge in biology, with the potential to unlock a wide range of applications, from the discovery of novel enzymes and therapeutic proteins to insights into evolutionary relationships and disease mechanisms^1–6^. This is particularly relevant for complex protein classes such as antibodies, nanobodies, and increasingly, *de novo* designed binders, which play indispensable roles in diagnostics, therapeutics, and biomedical research^7–13^. Their high specificity and affinity for target antigens enable applications ranging from cancer therapies^14^ to viral neutralisation^15^ and biomarker discovery^16^. Many antibodies are still generated through animal immunization or selected from (polyclonal) antibody libraries, where it is crucial to have a direct link between their genotype and phenotype (e.g., using B-cell sequencing or display technologies)^17–19^ to obtain their accurate sequence if the antibodies are to be recombinantly expressed^20^. Often, this entails cumbersome processes of handling living cells from animals or human donors, constructing antibody libraries, and/or access to costly equipment and highly trained scientists, which creates large barriers for many scientists that wish to sequence their antibodies, especially in the reagents field^17^. Additionally, antibody sequencing can be pivotal for mapping immune responses, guiding vaccine development, and supporting precision medicine approaches requiring detailed immune profiling^21–23^.

In addition to antibodies, reliable protein sequencing can find utility for sequencing of novel or engineered proteins for biothreat detection and biosafety, as advancements in generative protein design technology heighten potential risks associated with engineered proteins^24,25^. Rapid sequencing of proteins would enable early detection and intervention, allowing safety measures to be put in place faster. Beyond these applications, direct protein sequencing also holds promise in broader biomedical research, such as biomarker and enzyme discovery^26,27^ and mapping of protein-protein interaction networks^28^.

In contrast to nucleic acids, proteins cannot be readily amplified by polymerase chain reaction (PCR). As a result, while trace amounts of DNA or RNA are sufficient for high-throughput sequencing, protein sequencing requires significantly larger sample sizes. Currently, bottom-up mass spectrometry (MS), involving proteolytic digestion followed by liquid chromatography-tandem mass spectrometry (LC-MS/MS)^29^, is commonly employed for protein sequencing^4,30,31^. However, these approaches depend heavily on existing reference spectra^32,33^, limiting their utility for highly variable regions, like the Complementarity Determining Regions (CDRs) of antibodies, and are not applicable to novel proteins.

To address these limitations, strategies such as open searches and multi-protease digestion have been introduced to increase flexibility and sequence coverage. Open searches in proteomics allow identification of peptides without predefined post-translational modifications (PTMs)^34^ or protease cleavage specificity^35,36^, but the broad search space introduces significant false discovery rate (FDR) control challenges, increased computational costs, and is still dependent on reference databases. Similarly, utilizing multiple proteases for peptide generation enhances sequence coverage but complicates sample preparation and downstream analyses due to diverse cleavage specificities and increased methodological complexity^37,38^.

To circumvent these limitations, *de novo* peptide sequencing, which operates independently of reference sequence data, has emerged as a promising approach. Recent computational advancements, primarily through introduction of deep learning models, have significantly improved accuracy^39–41^. Nonetheless, critical challenges persist, including false-positive identifications, limited diversity in enzymatic digestion, sensitivity of assembly algorithms to sequencing errors, and the absence of robust workflows for peptide assembly into contiguous protein sequences.

In this study, we address the challenges associated with protein sequencing using mass spectrometry by introducing an enhanced methodology integrating optimized sample preparation protocols and advanced computational techniques enabling unprecedented accuracy in protein sequence determination. We use a complementary multi-enzyme digestion strategy that increases peptide diversity and generates overlapping ends, while maintaining simple and fast sample preparation. We utilize artificial intelligence-driven *de novo* peptide sequencing with rigorous post-processing and filtering of predicted peptides, reducing the number of false positives and facilitating accurate protein sequence assembly. Finally, we implement a protein sequence assembly method optimized to different protein modalities, along with alignment and clustering steps to refine and prioritise unique candidate sequences that span contiguous regions of target proteins. This integrated approach establishes a robust, template-agnostic platform for direct protein sequencing, with significant implications for the discovery of biomarkers, novel enzymes and biologics, as well as biothreat detection. Our direct protein sequencing workflow can be used to sequence any protein given enough purity, further expanding its applicability across diverse biological and industrial applications.

## Methods

### Samples

We used four different sample types to develop, optimise, and evaluate our methodology. These ranged in complexity and biological origin. Recombinant bovine serum albumin (BSA) was obtained by (Sigma, A9418).

The nanobodies (hereafter named Nb1-10) included in this study were discovered using phage display technology as described previously^42^. A list of the nanobodies alongside the methods used for their expression and purification has been described previously^41^.

The antibodies (hereafter named mAbB1-mAb5) included in this study are a wild-type human immunoglobulin (hIg) G1 antibody (2554_01_D11)^43^, three hIgG1 antibodies with LALA^44^ and YTE^45^ mutations (TPL0179_01_C08^46^, TPL0552_02_A05^47^, and TPL0039_05_A03)^47^, and an hIgG1 with LALA mutation (2555_01_A01)^43^. All mAbs were discovered using phage display technology, reformatted from an scFv to a full length IgG format and expressed in CHO cells followed by protein A purification as described previously^43^.

The three *de novo* mini-binders (miBds) (NY1-B04, SILSY1-G05, and SILSY1-B11) included in this study (hereafter named miBd1-3) were designed to bind the shared cancer antigens presented on the peptide major histocompatibility complex (pMHC). A detailed outline of the design process, as well as biophysical and functional characterisation has been published previously^48^. In brief, NY1-B04 was designed and identified to bind the pMHC HLA-A*02:01 presenting the peptide ‘SLLMWITQC’, while SILSY1-G05 and SILSY1-B11 are capable of binding to the neo-cancer antigen peptide ‘RVTDESILSY’ presented on HLA-A*01:01.

The miBds have been expressed in *Bacillus subtilis* A164Δ5 derivative^48^ and purified employing 6xHis-tag affinity purification. Briefly, the amino acid sequence of the miBd designs were complimented with an N-terminal secretion signal peptide from Savinase (MKKPLGKIVASTALLISVAFSSSIASA) and a C-terminal linking region (‘GGGGSEAAAKGGGGS’), in the case SILSY1-G05 and SILSY1-B11, followed by an Avitag (‘GLNDIFEAQKIEWHE’) and a 6xHis-tag (‘HHHHHH’). For NY1-B04, the linking region was omitted. The miBds were ordered as synthetic gene fragments and cloned in the expression vector pDG268Δneo. A recombinant clone containing the integrated expression construct was identified and grown in liquid culture medium.

For miBd purification, 250 mL of the sample containing cultures were subjected to centrifugation (10,000 x g for 15 min., Sorvall RC 6 plus, 46915 ThermoFisher Scientific), and the supernatant applied to single-use columns for immobilized metal affinity chromatography (His GraviTrap columns, 11003399, Cytiva) following the supplier’s protocol. Subsequently, the samples were desalted employing disposable PD-10 columns packed with Sephadex G-25 resin (17085101, Cytiva) and eluted in 5 mL of buffer solution (50 mM Tris, 150 mM NaCl, pH 7.4).

### Sample preparation for mass spectrometry

#### Bovine Serum Albumin (BSA)

The digestion protocol was optimized using bovine serum albumin (BSA) obtained in lyophilised form (Sigma-Aldrich, B6917). BSA was reconstituted in a sodium deoxycholate (SDC) lysis buffer containing 1% SDC, 200 mM Tris-HCl (pH 8.5), 10 mM Tris(2-carboxyethyl) phosphine hydrochloride (TCEP), and 40 mM 2-chloroacetamide (CAA). For each enzyme and condition, 10 µg of protein was transferred to Eppendorf Protein LoBind tubes and heated at 95°C for 5 minutes to ensure complete denaturation. Samples were then diluted 1:1 with 100 mM ammonium bicarbonate (pH 8.5) prior to enzyme addition. We tested a range of enzyme-to-protein ratios (1:50, 1:100, and 1:200), incubation times (1 hour, 4 hours, and 18 hours), and temperatures (23 °C and 37 °C) for the following proteases: papain (Sigma-Aldrich, P5306-25MG), proteinase K (Promega, V3021), chymotrypsin (Promega, V1061), GluC (Promega, V1651), thermolysin (Promega, P1512), and elastase (Sigma-Aldrich, 324681-50UG). Thermolysin was tested at 37 °C and 70 °C, and not at 23 °C. For thermolysin, chymotrypsin, and proteinase K, CaCl_2_ was added to a final concentration of 10 mM to act as a cofactor. Trypsin (Promega, V5280) and LysC (Wako, 125-05061) were only tested at 1:50 and 1:100 enzyme-to-protein ratio at 37 °C at the three incubation times.

Digestion was quenched by a 1:1 dilution with 2% trifluoroacetic acid (TFA), after which the samples were centrifuged at 14,000 x g for 20 min. to pellet the precipitated SDC. The resulting supernatants were desalted using SOLAµ SPE plates (Thermo Scientific, 60209-001), with all steps driven by centrifugation at 350 x g. The filters were activated with 200 µL of 100% methanol followed by 200 µL of 80% acetonitrile containing 0.1% formic acid. Columns were equilibrated twice with 200 µL of 1% TFA in 3% acetonitrile before sample loading. After loading, peptides were washed twice with 0.1% formic acid and subsequently eluted into clean 0.5 mL Eppendorf tubes using 40% acetonitrile with 0.1% formic acid. Eluates were concentrated in a SpeedVac (Eppendorf) and reconstituted in 12 µL of buffer A* (2% acetonitrile, 1% TFA) for peptide quantification by NanoDrop (DeNovix).

In addition to the enzymes with optimal activity at neutral pH, we also tested the performance of pepsin (Promega, V1959), legumain (Jena Bioscience, PR-967S), and Krakatoa (CinderBio, CB23726), which require acidic conditions. For these, BSA was reconstituted in a buffer containing 20 mM potassium phosphate and 20 mM citric acid, adjusted to pH 3 for pepsin and Krakatoa, and to pH 5.5 for legumain. TCEP and CAA were added to final concentrations of 10 mM and 40 mM, respectively, and the protein was denatured by heating to 95 °C for 5 minutes. Krakatoa was tested at a ratio of 1U enzyme per 5 µg protein (according to the supplier’s instructions) and incubated at 80 °C for 5, 30, and 60 minutes. Pepsin and legumain were evaluated under the same conditions used for the neutral pH enzymes. All reactions were quenched by placing the tubes on ice, and peptide clean-up and reconstitution was performed as described above.

After reconstitution in A* buffer, 50 ng of each sample was analyzed on LC-MS. Data was analyzed to find conditions, which result in high numbers of peptides while simplifying sample preparation, to avoid overly complex experimental setup. The resulting conditions were used for the processing of Nbs, mAbs, and miBds.

#### Nanobodies

A total of 10 µg of each Nb for each enzyme digest was diluted with SDC lysis buffer to a final volume of 100 µL. CaCl_2_was added to a final concentration of 10 mM to the SDC lysis buffer used for samples that were to be digested with chymotrypsin, proteinase K or thermolysin. Samples were heated to 95 °C for 5 minutes and diluted 1:1 with 100 mM ammonium bicarbonate (AMBIC), pH 8.5. Enzymes were added in the following enzyme-to-protein ratios; papain, GluC, and elastase 1:50, trypsin 1:100, and chymotrypsin, proteinase K, and thermolysin 1:200. All samples were incubated at 37 °C for 4 hours, except the proteinase K digests, which were acidified after 1 hour. Samples were acidified by diluting 1:1 with TFA 2%. Samples were centrifuged at 14,000 x g for 20 minutes to pellet the precipitated SDC. Additionally, nanobodies were digested using Krakatoa (CinderBio, CB23726) and Vesuvius (Cinderbio, CB14057). Here, 10 µg of the nanobodies were mixed with TCEP and the provided digestion buffer according to the supplier’s instructions. pH was confirmed to be 3 and enzyme was added in a 2U/µg sample ratio. Samples were incubated at 80 °C for 30 minutes and put on ice to inactivate the enzymes. Supernatants containing resulting peptides were desalted and reconstituted using the above mentioned procedure. A total amount of 100 ng of peptides from each sample was injected and analyzed with LC-MS/MS.

#### Antibodies

A total of 10 µg of each antibody or antibody mix for each enzyme digest was diluted with SDC lysis buffer to a final volume of 100 µL. Samples were heated to 95 °C for 5 minutes and diluted 1:1 with 100 mM AMBIC pH 8.5. Calcium chloride was added to a final concentration of 10 mM to the mAbs for digestion with thermolysin, proteinase K and chymotrypsin. The proteases papain, GluC, elastase, trypsin, chymotrypsin, proteinase K and thermolysin were added to the respective samples following the same ratios, incubation times and temperatures as mentioned in section 3.2.2. TFA was added to a final concentration of 1% to stop enzyme activity and precipitate SDC, and samples were centrifuged at 14000 x g for 20 minutes. Supernatants were desalted and reconstituted following the same procedure as above. 100 ng of each sample was analyzed with LC-MS/MS.

#### Mini-binders

MiBds were prepared for MS analysis with the exact same protocol as nanobodies.

### MS acquisition

Peptides were loaded for analysis onto a 2 cm C18 trap column (Thermo Scientific 164946), connected in-line to a 15 cm C18 reversed-phase analytical column (Thermo EasySpray ES904) using 100% Buffer A (0.1% formic acid in water) at 750 bar, on the Thermo EasyLC 1200 HPLC system, with the column oven set to 30 °C.

Peptides were eluted using a 35-minute gradient at a flow rate of 250 nL/min. The gradient began with a transition from 10% to 23% Buffer B (80% acetonitrile, 0.1% formic acid) over 17 minutes and then increased to 38% Buffer B over 6 minutes. This was followed by ramping up to 60% Buffer B of 3 minutes, after which Buffer B was increased to 95% over 3 minutes, holding at this concentration for 6 minutes.

The Q-Exactive instrument (Thermo Scientific) was run in a DD-MS2 top 10 method. Full MS spectra were collected at a resolution of 70,000, with an AGC target of 3×10 e^6^ or a maximum injection time of 20 ms and a scan range of 300–1750 m/z. The MS2 spectra were obtained at a resolution of 17,500, with an AGC target value of 1×10 e^6^ or a maximum injection time of 60 ms, a normalized collision energy of 25, and an intensity threshold of 1.7×10 e^4^. Dynamic exclusion was set to 30 s, and ions with a charge state 1, >6 or unknown were excluded. MS performance was evaluated by running complex cell lysate quality control standards.

### Database search

Raw files were analyzed using Proteome Discoverer 2.4 (Thermo Scientific) software. The LC-MS/MS raw files were processed in batches based on the enzyme used for cleavage and with the use of label-free quantitation (LFQ) in both the processing and consensus steps. In the processing step, oxidation (M), and protein N-termini acetylation and met-loss were set as dynamic modifications, with cysteine carbamidomethyl set as static modification. All results were filtered with Percolator using a 1% false discovery rate (FDR), and Minora Feature Detector was used for quantitation. SequestHT was used as a search engine, matching spectra against the uniprot bovine database for the BSA samples, contaminant database, and known mAb sequences. Nbs and miBds were analysed directly with *de novo* sequencing.

### *De novo* peptide sequencing

We performed *de novo* peptide sequencing using InstaNovo v0.1.4^41^. We first converted raw mass spectrometry (MS) data into the standardized .mzML format using ProteoWizard MSConvert^49^. We put the resulting .mzML files into the InstaNovo framework, which we deployed on a high-performance computing (HPC) cluster (https://www.hpc.dtu.dk/). Results were exported and used for downstream analysis with InstaNexus.

### Protein sequencing and assembly

#### Data processing

The data processing steps after *de novo* peptide inferencing include the following key steps in the order they are presented: We converted InstaNovo’s logarithmic confidence score to its exponent for improved interpretability, obtaining a confidence score from 0-1 for each peptide prediction. We stripped modification strings included in the peptide sequences (e.g., (ox)) to standardise formatting and enable sequence mapping. We discarded rows that contain missing values in predictions, as these were very low confidence predictions removed already by the tool. We selected high-confidence peptide-spectrum matches (PSMs) to reduce the number of false positives during assembly. We also optimized to obtain a balance of predictions that map to our targets, and false positives or predictions that were peptides from other proteins present in the sample. Finally, we excluded predictions that map to known contaminants, such as albumin and collagen sequences.

#### Greedy peptide assembly

We performed peptide assembly using two complementary strategies: a greedy overlap-based method and a de Bruijn graph (DBG) approach, each designed to maximise sequence reconstruction based on the predictions generated by InstaNovo.

The greedy assembler operates by identifying complete overlaps between PSMs, specifically where the suffix of one peptide aligns with the prefix of another. Sequences are iteratively merged based on a defined minimum overlap threshold, continuing until no further integration is possible. To reduce redundancy, which is especially relevant due to the use of multiple proteases, assembled contigs were filtered to eliminate duplicates and overlapping sequences. Starting from these initial contigs, a scaffolding step was applied, which progressively extends the sequences by increasing the required minimum overlap, improving contig length and continuity. We consider it a critical methodological choice to set the minimum overlap to at least 3 amino acids to prevent erroneous contig generation.

#### Peptide assembly with de Bruijn graphs

In contrast to greedy assembly, the de Bruijn graph (DBG) method fragments each peptide sequence into overlapping subsequences, or k-mers. Each k-mer represents a node in the graph, and edges are defined by overlaps of k−1 amino acids between adjacent k-mers. Assembly proceeds through graph traversal using a depth-first strategy to identify valid paths that assemble longer contiguous sequences. During this traversal, additional filters are applied to remove duplicates, enforce minimum scaffold lengths (we use the term scaffolds to indicate merged contigs derived from greedy or DBG assembly), and prioritise sequence uniqueness.

For the DBG-based method, the optimized parameters included: 1) the minimum confidence threshold for accepting PSMs; 2) the k-mer size used for graph construction; 3) the minimum overlap required to merge contigs into scaffolds; 4) the maximum number of mismatches and the minimum alignment identity for mapping contigs to the reference protein sequence; and 5) the minimum sequence length threshold to retain valid contigs or scaffolds. While k-mer size is a key parameter in de Bruijn graph-based approaches, it was fixed at 7 in this study, since for all sample types similar performance was observed for k-mer sizes of 6 or 7, with dramatically reduced performance with any other size.

#### Clustering, alignment, and consensus sequence generation

We clustered assembled scaffolds using MMseqs2^50^ with a minimum identity threshold set to 0.85 and a coverage mode of 1 to merge near-identical sequences and reduce redundancy arising from overlapping or partially assembled contigs. A minimum identity threshold of 0.85 ensures that only sequences sharing at least 85% amino acid identity over the aligned region are grouped into the same cluster, thus preserving sequence specificity. With coverage mode 1, MMseqs2 computes the alignment coverage as the fraction of the target sequence that is aligned to the query:

Coverage = Number of aligned residues / Length of target sequence

For each resulting cluster, we performed multiple sequence alignment using Clustal Omega version 1.2.4^51^, with default settings. From each aligned cluster, we computed a position-specific scoring matrix (PSSM) by calculating the relative frequency of each amino acid at every aligned position, excluding gaps. This matrix is subsequently used to derive a representative consensus sequence. At each aligned position, we selected the amino acid with the highest frequency among the aligned sequences.

#### Parameter optimization and assembly evaluation

To optimize the performance of the assembly algorithm, a systematic hyperparameter search was carried out using both grid search and de Bruijn graph strategies. A structured set of parameters – including confidence threshold, scaffold overlap thresholds, maximum mismatches, minimum overlap, minimum alignment identity, and minimum size thresholds – was explored by generating all possible combinations across predefined ranges. This exhaustive exploration produced 1,280 distinct configurations for each sample. To efficiently manage the computational workload, the search was executed in parallel using Python’s ProcessPoolExecutor, leveraging up to 64 workers (threads) concurrently. The primary goal of this optimization was to maximise assembly coverage. However, coverage alone is not a sufficient criterion. At equal levels of coverage, other metrics were taken into account, such as the total number of assembled sequences, maximum scaffold length, and N50/N90 values.

#### Composite score

To comprehensively assess assembly quality, we computed a composite score that integrates multiple complementary metrics. Each metric was first normalized individually using MinMaxScaler, which linearly scales the original values to a range between 0 and 1. This normalisation step ensures that all metrics contribute comparably to the final score, regardless of their original scale or units. The composite score for each assembly was calculated as the dot product between the vector of normalized metrics and the corresponding vector of weights, following the formula:

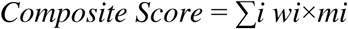

where *wi* is the weight assigned to metric *i*, and *mi* is the normalized value of that metric. Assemblies were subsequently ranked according to their composite scores to prioritise those demonstrating the best overall balance of coverage, contiguity, fragmentation, and completeness. We assigned weights based on the relative importance of each metric in reflecting key aspects of assembly quality. Coverage received the highest weight (0.5), as it represents the most critical indicator of the extent to which the input data support the assembled sequences. N50 was assigned a weight of 0.3, capturing contiguity and reflecting the ability of the assembly to produce longer and coherent sequences. The total number of sequences was weighted at 0.1. Since fewer sequences typically indicate reduced fragmentation and improved assembly quality, we inverted the normalized values (i.e., computed 1 minus the normalized value) to ensure that assemblies with fewer fragments received higher scores. Finally, the maximum length, also weighted at 0.1, was included to favor assemblies capable of recovering complete or near-complete protein regions.

Pipeline reusability and repository structure

The InstaNexus workflow was implemented using Jupyter Notebooks, accessible in this GitHub repository https://github.com/Multiomics-Analytics-Group/InstaNexus. The repository contains the core analysis scripts and auxiliary functions needed to reproduce the results. The repository follows a standard Python package structure with a “src” directory containing the Python scripts required for executing the analysis, including the hyperparameter optimization module, the “notebooks” directory with the Notebook files that can be reused to run the greedy and DBG methods, and the “analysis” directory that includes secondary analyses and plotting functionality. The Notebook design allows users to independently modify or extend the workflow. All dependencies and instructions are documented in the repository’s README files.

## Results

### InstaNexus, an optimized workflow for direct protein sequencing

Our direct protein sequencing workflow InstaNexus starts with a multi-protease panel for enzymatic digestion prior to MS analysis (Figure 1A). We included proteases covering a broad specificity profile to ensure the generation of diverse and overlapping peptides and to maximise compatibility with a broad sample repertoire. We tested 14 different proteases and optimized reaction conditions and incubation times individually for each protease. Thereafter, the performance was evaluated by the number of unique peptides detected through database searches using BSA as a model sample. Our optimized sample preparation workflow includes 10 proteases, each digesting the sample in separate reactions. The sample preparation is completed in less than four hours, making the process highly time- and cost-efficient. Moreover, only 5-10 µg of material is required per protease digestion, thus minimizing the sample amount needed for analysis. For peptide prediction, we selected InstaNovo as the foundation for *de novo* sequencing due to its high performance across a wide range of biological contexts^41^. InstaNovo delivers accurate and generalizable peptide identifications, including non-tryptic peptides and modified residues, which are essential for robust downstream assembly (Figure 1B).

**Figure 1.**
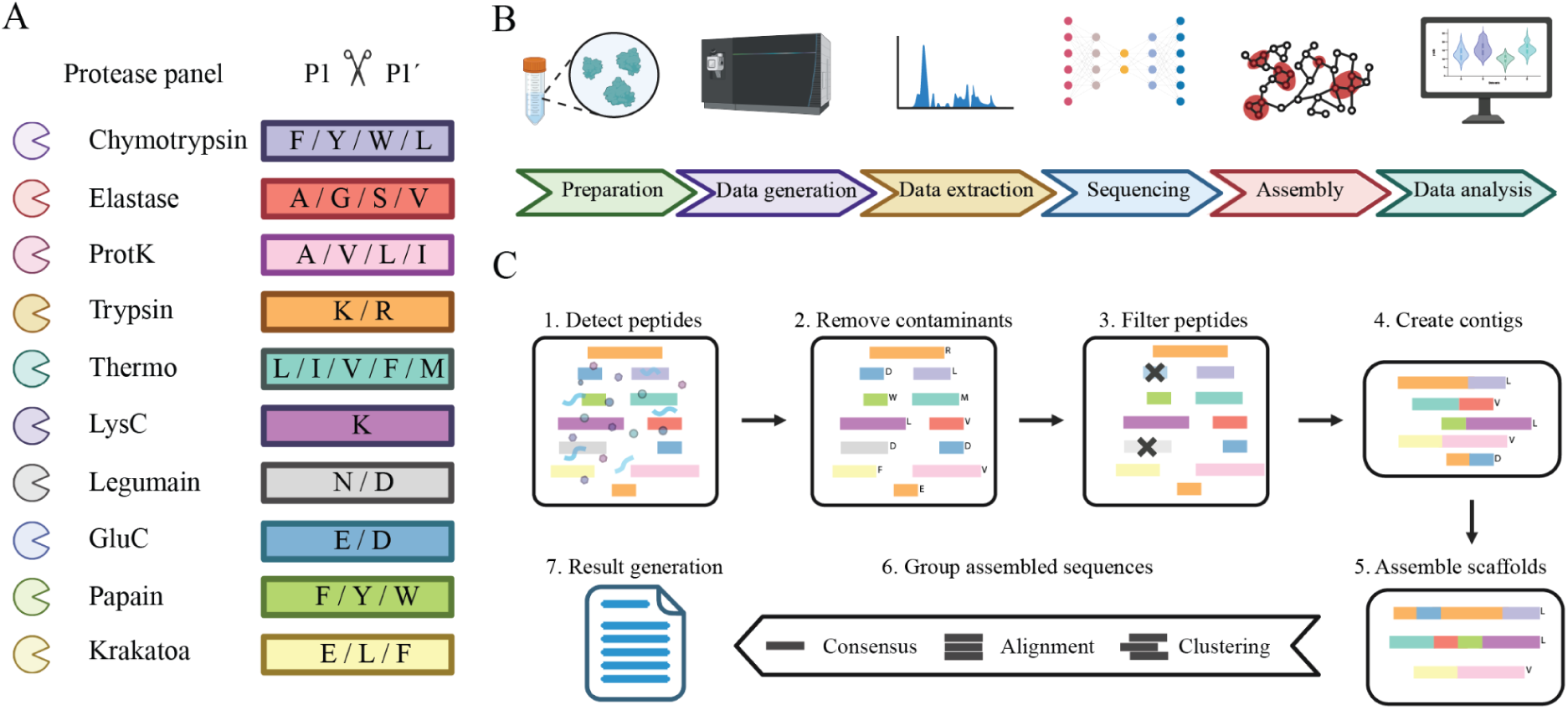
Overview of the optimized direct protein sequencing workflow. (A) A multi-protease panel covering a broad cleavage specificity range was established for enzymatic digestion. Proteases were selected and optimized to generate overlapping peptides, maximizing sequence coverage and reducing sample requirements. (B) Overview of the experimental and computational workflow: sample preparation data generation via mass spectrometry, peptide extraction, de novo sequencing using InstaNovo, assembly, and final data analysis. (C) Assembly computational pipeline: (1) removing contaminants, (2) cleaned de novo peptide predictions, (3) filtering of low confidence peptides, (4) assembly into contigs using greedy or DBG approaches, (5) scaffold formation, (6) alignment, clustering and consensus generation, resulting in (7) a final set of candidate protein sequences.

InstaNexus assembles protein sequences using two complementary computational methodologies: a greedy assembler and a DBG-based assembler. These approaches independently generate overlapping peptide contigs, which can then be merged to produce longer sequence scaffolds, the term we use to describe meta-contigs or assembled contigs following the terminology in genomics. In the early stages of assembly, duplicates or fully contained contigs within longer ones are removed to reduce complexity. The remaining non-redundant contigs are then merged into scaffolds to further extend sequence length. The resulting scaffold space represents the candidate protein sequences. However, several scaffolds may still be partially overlapping or contain positional disagreements. To consolidate this space, we align scaffolds using ClustalOmega and perform clustering with MMseqs2. From each cluster, we generate a frequency matrix and derive a consensus sequence, reducing the scaffold pool to a minimal, high-confidence set of candidate sequences that cover the target sequence with high information content (Figure 1C).

### InstaNexus generates high quality predictions and sequence coverage at peptide level

To benchmark and optimize the computational components of our workflow, we first applied InstaNexus to BSA, a standard protein with a well-characterized reference sequence. InstaNovo generates a large number of peptides with assigned confidence scores capturing the probability of a correct peptide identification. The output peptides included a subset of peptide-spectrum matches (PSMs) without assigned confidence scores (15,709 out of 109,358; 14.36%) and further filtering was required to ensure data quality and accurate downstream analysis (Supplementary Figure 1). The correlation between confidence scores and peptide identification accuracy highlighted InstaNovo’s ability to prioritise high-quality peptide candidates (Figure 2A). To define an optimal confidence threshold, we quantified the proportion of mapped versus unmapped peptides across increasing confidence intervals (Figure 2B). We established a cutoff where the number of mapped peptides exceeded the number of unmapped peptides, filtering out the majority of false positive identifications and enhancing the precision of downstream sequence reconstruction without sacrificing sequence coverage (Figure 2C). On BSA, this threshold corresponded to a confidence score of 0.88, where the mapped/unmapped peptide ratio reached approximately 1:1 (Figure 2D). This threshold was used as a reference point for subsequent analyses. Particularly for less complex samples or recombinant proteins, we found that a confidence level near 90% was required to achieve a mapped/unmapped ratio greater than 1.

**Figure 2.**
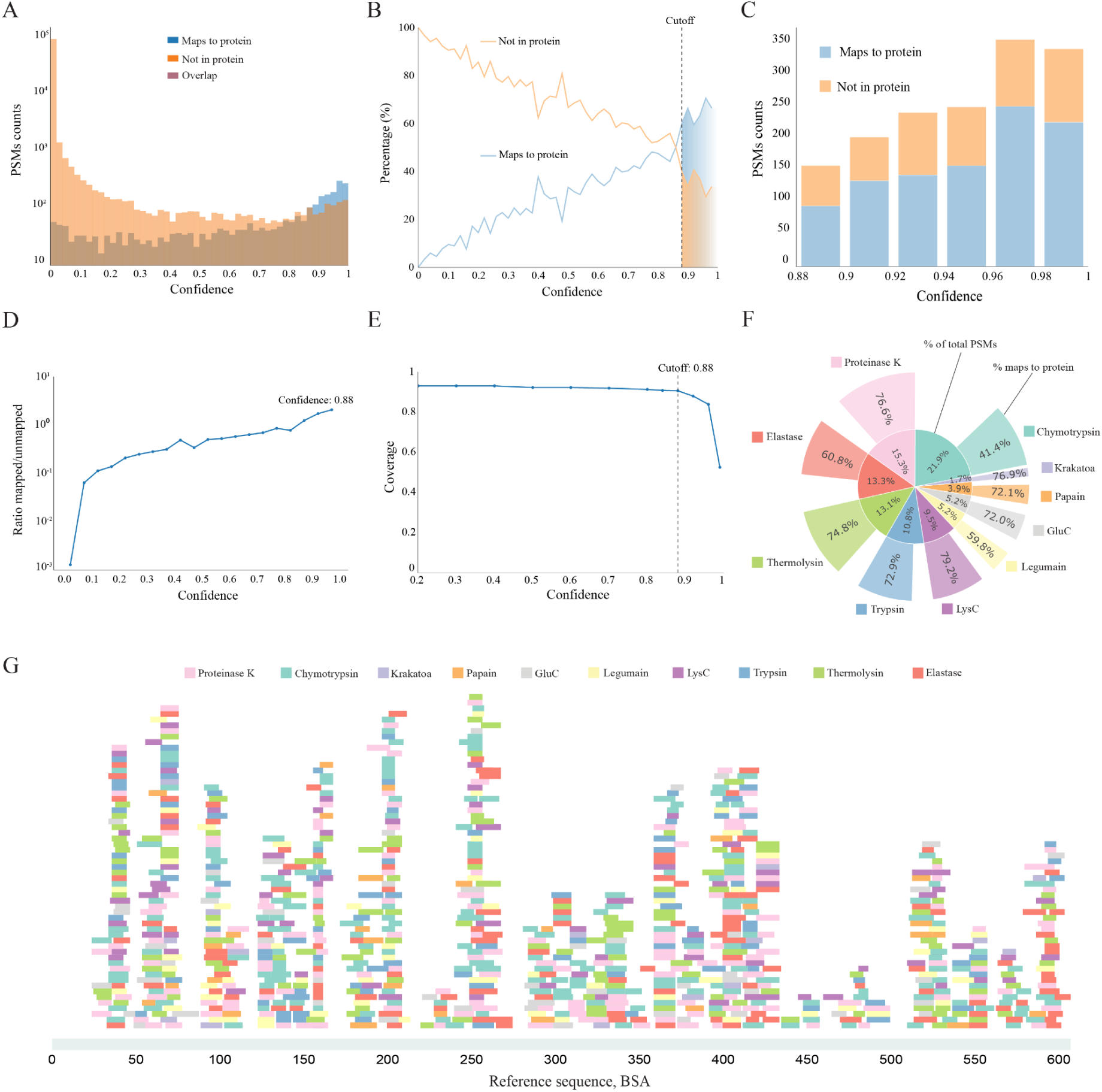
Evaluation of peptide prediction quality and sequence coverage. (A) Distribution of peptide-spectrum matches (PSMs) across confidence scores, showing mapped peptides (blue), unmapped peptides (orange), and overlap (brown). (B) Percentage of mapped and unmapped peptides across the confidence range. A confidence cutoff of 0.88 is selected where mapped peptides surpass unmapped ones. (C) Distribution of PSMs above different confidence thresholds, categorized by mapping status. (D) Ratio of mapped to unmapped peptides as a function of confidence score. A threshold around 0.88 maximises enrichment for true positives. (E) Sequence coverage by mapped peptides remains high until the 0.88 cutoff, then drops with stricter filtering. (F) Contribution of each protease to the total PSMs and mapping efficiency to the reference sequence. The inner segments represent the percentage of total PSMs attributed to each protease, while the outer segments reflect the fraction of those PSMs that successfully map to the reference protein sequence. (G) Mapping of peptides along the bovine serum albumin reference sequence, with each protease contributing complementary coverage.

We next assessed the impact of confidence filtering on sequence coverage. Coverage remained saturated up to the confidence score of 0.88 threshold, followed by a steep decline at high stringency levels (Figure 2E). Notably, high-confidence peptides that mapped to the protein target maintained coverage of the protein sequence while reducing contributions from low-confidence predictions, potential contaminants or other host cell proteins in the sample. These findings confirmed that this confidence threshold effectively maximises true positive contigs and preserve near-complete sequence coverage.

To evaluate the contribution of individual proteases to peptide detection and mapping accuracy, we analyzed their performance in terms of both total PSM yield and mapping efficiency (Figure 2F). Proteases such as trypsin, LysC, thermolysin, and chymotrypsin contributed not only with a high proportion of total PSMs but also showed mapping rates exceeding 70%, indicating high efficiency in generating detectable peptides. In contrast, enzymes like proteinase K, Krakatoa, and papain yielded lower mapping percentages, which may reflect either broader cleavage patterns or reduced compatibility with InstaNovo, potentially leading to unmapped or ambiguously identified peptides (Supplementary Figure 2). Mapping of high-confidence peptides across the BSA sequence showed broad, complementary coverage resulting from combined protease-specific digestions (Figure 2G), underscoring the value of our multi-protease strategy.

### Hyperparameter tuning improves scaffold-level coverage by identifying optimal settings

After establishing high coverage and accuracy at the peptide level, we sought to optimise the parameters of our multi-step assembly workflow to maximise candidate sequence quality (Figure 3A). Using our BSA dataset, we systematically evaluated sequence coverage resulting from different parameter combinations using a grid search approach. After testing optimal values for six parameters, we visualized the impact of four key parameters: confidence threshold, size threshold, minimum identity, and maximum mismatches (Supplementary Figure 3). Although the assembly process yielded multiple contigs, which correlated with the size of BSA, we achieved deep coverage and accurate sequence assembly across the full-length protein (Figure 3B). Notably, the overall coverage would be even higher if the N-terminal signal peptide, which is typically cleaved in the mature form of BSA, was not included in the reference sequence. Mapping analysis also revealed mismatches and frequent amino acid substitutions, in particular N→D and Q→E deamidation events compared to the reference sequence. We constructed a frequency matrix from each cluster containing aligned scaffolds, which demonstrated high confidence and discrimination across positions (Figure 3C). From the PSSM, we derived a sequence logo plot to visualise the most likely amino acid candidates at each position within the generated sequence, which are in agreement with the reference sequence (Figure 3D).

**Figure 3.**
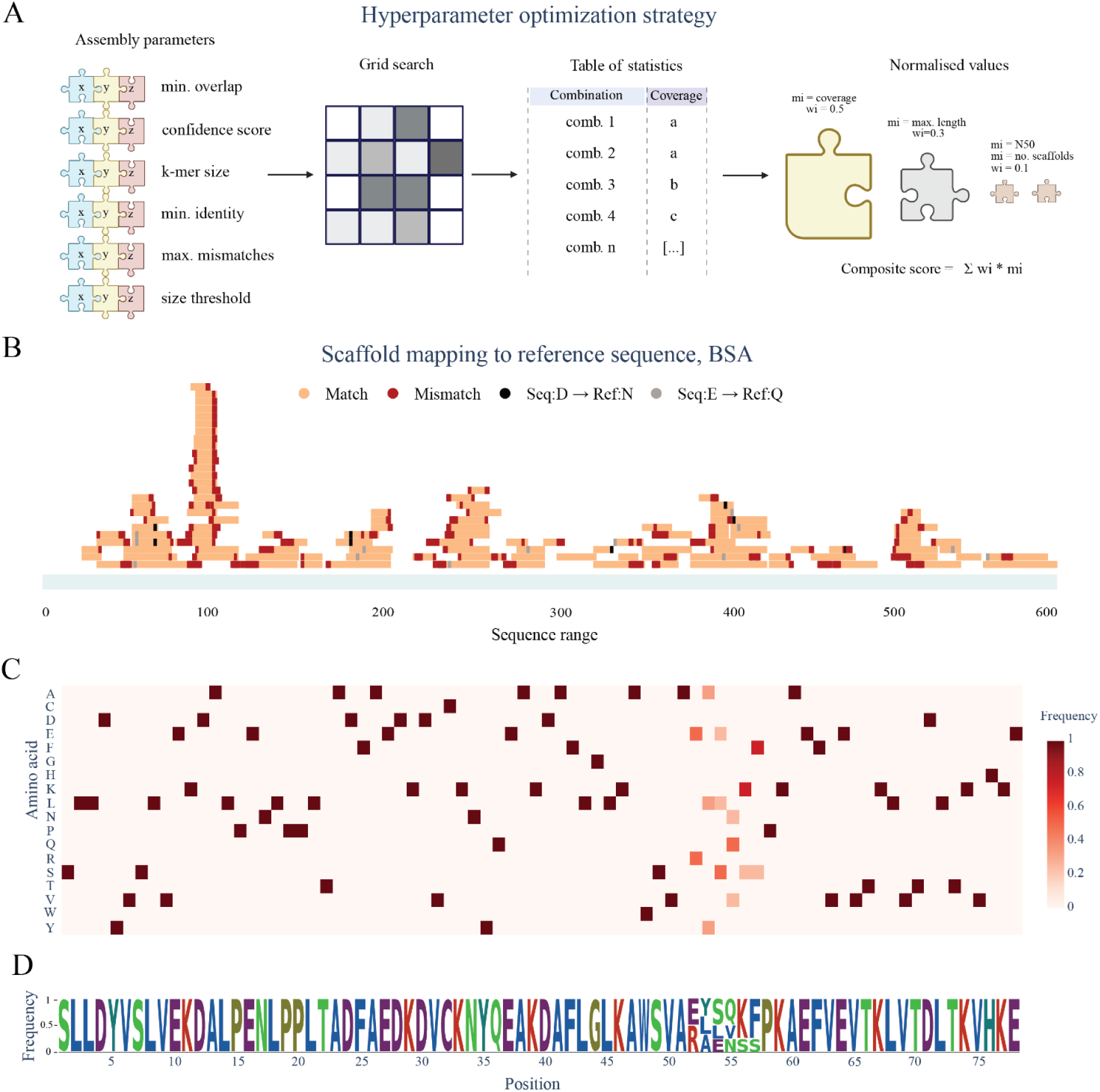
Parameter optimization and consensus-based scaffold assembly. (A) Hyperparameter optimization for scaffold assembly. Assembly parameters were varied in a grid search and combinations were scored using normalized metrics to identify optimal settings. (B) Mapping of assembled scaffolds to the bovine serum albumin reference sequence. Matches (light orange), mismatches (red), and known modification-driven substitutions (N→D, Q→E) are annotated. (C) Position-specific scoring matrix (PSSM) of amino acid frequencies across aligned scaffolds. (D) Sequence logo plot derived from the PSSM showing residue prevalence across the consensus.

### InstaNexus produces long contigs and detects hypervariable regions in nanobodies

Our method’s performance with BSA, suggests its potential for application to other proteins. Therefore, we set out to sequence a panel of 10 nanobodies using both contig and scaffold-level assemblies. Overall, we achieved high maximum coverage across the majority of nanobodies (Figure 4A), with comparable performance between contigs and scaffolds. We analyzed the PSM depth across the nanobody sequence, focusing on the CDRs (Figure 4B). The three CDRs exhibited distinct coverage patterns, with CDR2 showing the highest and most consistent signal. In contrast, CDR1 was preceded by a marked drop in PSM depth, while CDR3 coverage remained moderate. To evaluate the quality of the assemblies, we developed a composite score, incorporating downstream metrics such as number of final sequences and length. Composite scores ranged from 0.642 (Nb1) to 0.7 (Nb8), with nanobodies 7, 8, and 10 exhibiting the highest values (Figure 4C and Supplementary Table 1).

**Figure 4.**
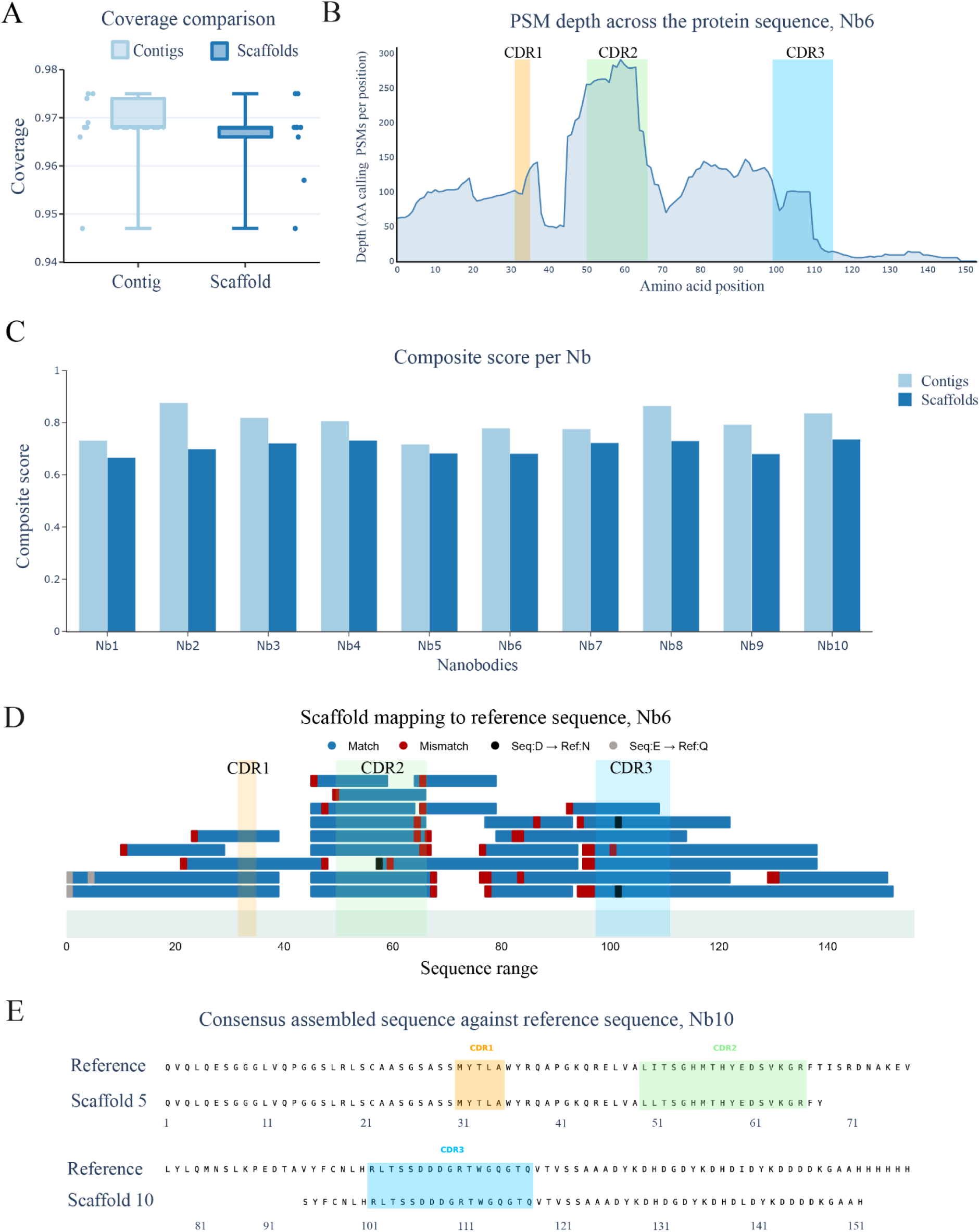
Sequence reconstruction accuracy across nanobodies. *(A)* Coverage comparison between contig-based (light blue) and scaffold-based (dark blue) assemblies across the sequences for the nanobodies included in this study. (B) PSM depth across the sequence for nanobody 6. The y-axis indicates the number of matching PSMs per amino acid position. All three complementarity-determining regions (CDR1-CDR3 shaded) are well covered, with CDR2 showing the highest signal intensity. (C) Composite scores per nanobody, comparing contig and scaffold-based assemblies. Contigs generally show improved scores across nanobodies. (D) Scaffold-to-reference sequence alignment for nanobody 6 showing matches (blue), mismatches (red), and sequence ambiguities (grey bars). (D) Consensus sequence comparison for nanobody 6, aligned to the reference. All three CDRs are accurately recovered. Notably, in scaffold 5, both CDR1 and CDR2 are recovered with maximal accuracy within a single scaffold.

We selected nanobody 6 for detailed analysis as a representative case. Scaffold-to-reference mapping revealed a largely accurate alignment across the sequence range (Figure 4D), with high match frequency interspersed by localized mismatches and evidence of deamidation events. These discrepancies were predominantly concentrated near the 5’ and 3’ termini, which also coincided with repetitive regions. Similar results were observed across nanobody 6 using the DBG method in Supplementary Figure 4, further supporting the high performance and consistency of the DBG-based approach.

To gain deeper insight into region accuracy at the functional level, we directly aligned the scaffold for Nb10 against the reference sequence. Importantly, we were able to recover all three CDRs with complete accuracy (Figure 4E).

### An accurate workflow for antibody sequencing

We next assessed the performance of our workflow by sequencing three mAbs. High maximum coverage values were consistently observed across all mAbs, with comparable performance between contigs and scaffolds (Figure 5A). Composite scores indicated robust sequence accuracy across mAbs, with mAb2 showing slightly higher composite scores compared to others (Figure 5B). For detailed analysis, we selected the heavy chain of mAb1. Scaffold-to-reference mapping indicated accurate sequence reconstruction predominantly towards the C-terminal region of the protein (Figure 5C), characterized by frequent matches interspersed with sporadic mismatches. Typical deamidation events were also observed as previously. To further challenge our workflow, we pooled five distinct mAbs, including the three previously mentioned, and analyzed the resulting oligoclonal antibody mixture. To evaluate optimal parameters and maximum performance when the reference sequence was known, each mAb underwent a systematic optimization of assembly parameters. The composite score was then used to identify the optimal assembly conditions for each antibody (mAb1, mAb2, and mAb3). The average of these optimized parameters was subsequently applied to the dataset for the oligoclonal antibody mixture, enabling a direct comparison of performance metrics (Supplementary Table 2) between monoclonal and oligoclonal sequence generation. Compared to the oligoclonal mixture, the mAb1 light chain showed consistently higher values across all metrics used in the composite score, such as N50 in the monoclonal sample, indicating superior assembly performance (Figure 5D). For the heavy chain, the monoclonal sample also outperformed the oligoclonal mixture, particularly in sequence coverage, composite score, and total sequences generated, although the difference in N50 was less pronounced. A detailed comparison of the mAbs within the oligoclonal mixture compared to their monoclonal sample counterparts, including metric-by-metric trends and variability across optimization conditions is provided in Supplementary Figure 5. For the light chain, sequencing of mAb2 showed similar performance in comparison to retrieving the same sequence in the oligoclonal mixture, with only minor gains in N50 and composite score. In contrast, its heavy chain assembly is clearly superior in coverage, N50, maximum length, and composite score. Surprisingly, mAb3 sequence assembly outperforms the oligoclonal mixture in the light chain for N50, max length, and composite score. In the heavy chain, however, the monoclonal sample consistently outperforms the assembly within the oligoclonal mixture across all statistics. Overall, monoclonal assemblies yield higher assembly metrics compared to the oligoclonal mixture, particularly for the heavy chains. This is not surprising, given the increasing sample complexity and ambiguity created when assembling proteins with mostly similar sequences.

**Figure 5.**
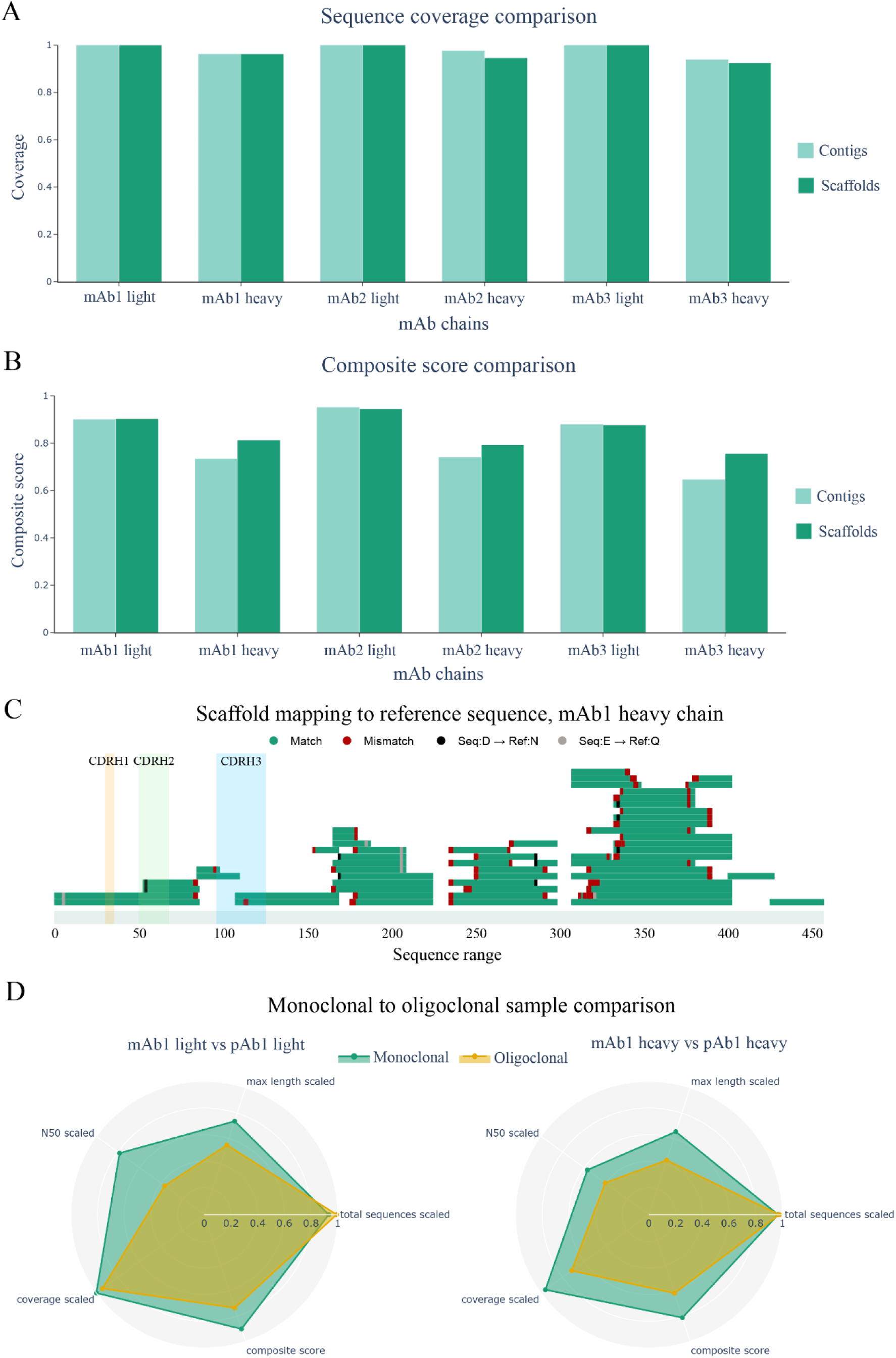
Assembly performance across monoclonal and oligoclonal mAbs. (A) Maximum coverage for each monoclonal antibody chain (heavy and light), measured for contigs (light green) and scaffolds (dark green). (B) Composite scores for all monoclonal antibody chains, comparing contig and scaffold-level assemblies. (C) Scaffold-to-reference mapping for the heavy chain of monoclonal antibody 1 (mAb1), indicating sequence matches (green), mismatches (red), and deamidation substitutions. (D) Scaled performance metrics for mAb1 sequenced individually versus as part of the oligoclonal antibody mixture, shown separately for the light and heavy chains. Metrics include total sequences, maximum length, N50, coverage, and composite score.

### InstaNexus can sequence de novo designed binders

Finally, we evaluated the capability of our workflow to reconstruct sequences of miBds. We analyzed three miBds designed *in silico* to target shared cancer antigens presented on major histocompatibility complex molecules. The overall maximum coverage observed for miBds was lower than for Nbs and mAbs. Nevertheless, miBd3 showed nearly 100% sequence coverage and high composite scores for contig and scaffold assemblies (Figure 6A). In contrast, miBd1 and miBd2 showed lower values across both metrics. Among the three miBds, miBd3 was selected as the representative candidate due to its highest coverage and composite score (Figure 6B and Supplementary Table 3). Mapping of assembled scaffolds revealed good coverage across the reference sequence, with a notable gap at position ∼40. The C-terminal region of the sequence showed the highest number of overlapping mapped scaffolds. However, we observed a higher rate of mismatches compared to other samples, especially near scaffold boundaries, while deamidation events were only observed in the C-terminal region (Figure 6C). We next examined the longest scaffold obtained for miBd3. Our consensus sequence visualized with a sequence logo plot derived from this scaffold confirmed recovery of a 68-residue stretch (Figure 6D). Structural modeling based on the recovered sequence showed a well-formed helical conformation, with backbone geometry closely matching the reference structure (Figure 6E). Mapping scaffold coverage onto the structure revealed that some regions were not sequenced, likely due to sequence features rather than structural differences. In particular, the C-terminal region included a 6xHis-tag, which may be underrepresented due to difficulties in assembling low-complexity or repetitive sequences. Similarly, around position 40, the sequence includes two short repeated stretches (AAARHVAAA), which might be filtered out under stringent thresholds for minimum identity or maximum allowed mismatches.

**Figure 6.**
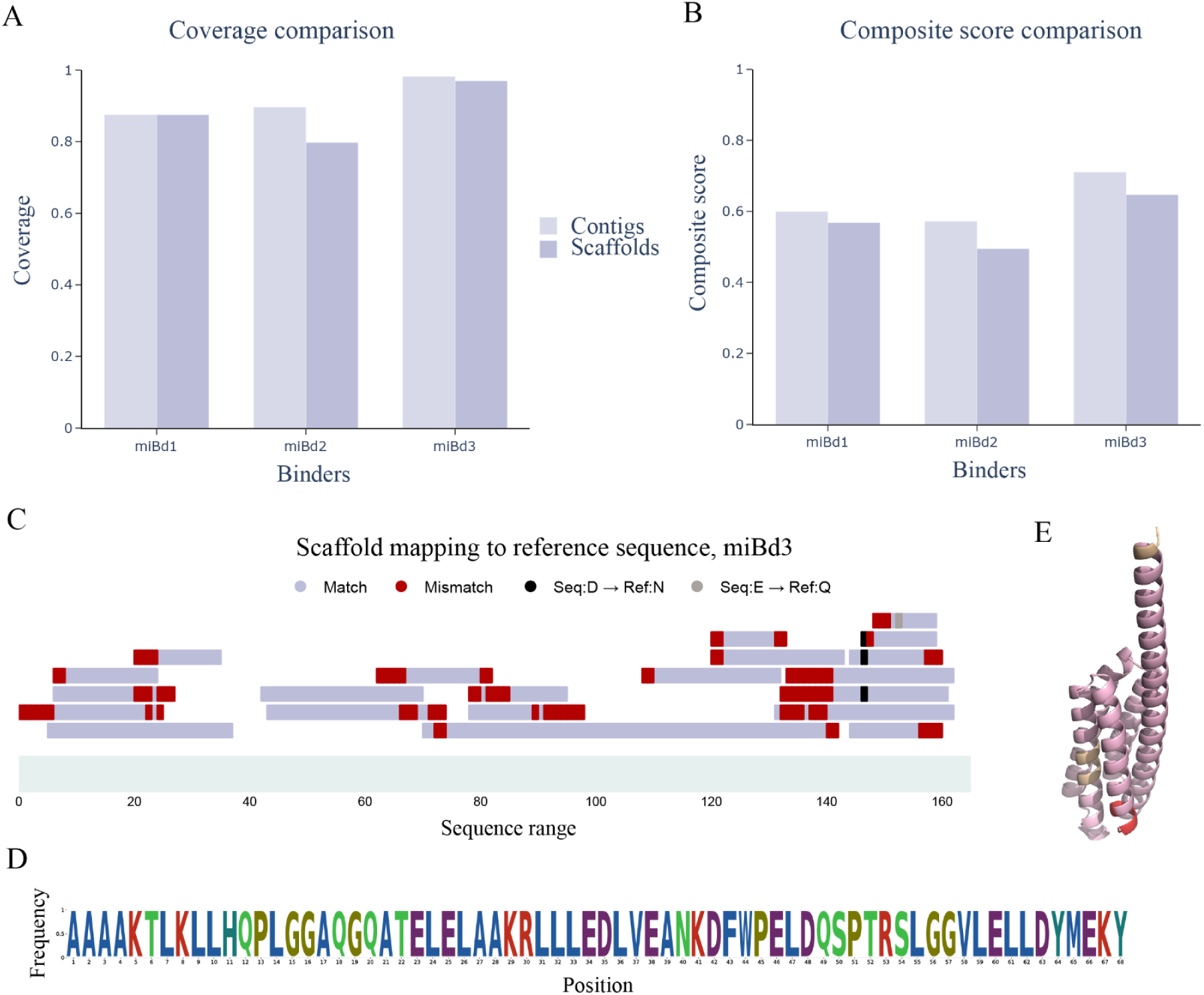
Sequencing performance for de novo designed miBds. (A) Maximum coverage and composite scores for three designed miBds, comparing contigs (light purple) and scaffolds (dark purple). (B) Composite scores for three designed miBds, comparing contigs and scaffolds. MiBd3 shows the highest performance coverage and composite score. (C) Mapping of scaffolds to the miBd3 reference sequence. Matches (dark purple) and mismatches (red) are shown along the sequence. Mapping is equally distributed across the protein. (D) Sequence logo plot derived from a scaffold of miBd3 covering the initial 68 residues. (E) Structural model of mapped scaffold coverage in miBd3: purple regions indicate positions covered by at least one scaffold (including those with mismatches), red highlights positions with mismatches only (no matching residues), and grey marks uncovered regions.

## Discussion

Despite the critical need for direct protein sequencing in diverse applications in biotechnology, current approaches face significant challenges including complex laboratory protocols and data acquisition, and require large sample amounts as well as heavy manual sequence inspection. In addition, there is inherent difficulty in generating sufficiently long and contiguous peptide sequences, and the overall absence of integrated, end-to-end workflows capable of managing this assembly complexity effectively. Our work presents an optimized workflow for direct protein sequencing, combining an improved laboratory protocol and a new bioinformatic pipeline to assemble protein sequences with near complete coverage with only a few contigs. Our sample preparation protocol leverages a diverse set of proteases to generate highly overlapping peptides, critically facilitating robust protein sequencing. A recent study used the proteases Krakatoa and Vesuvius to generate overlapping peptides for antibody sequencing and demonstrated higher accuracy compared to using single protease digestion, validating the selection of these proteases in our panel^38^. Crucially, our optimized sample preparation approach employs consistent buffer compositions wherever possible and limits digestion times to 4 hours. This enables high-throughput preparation of samples, automation of nearly all steps before MS analysis, and result generation within a single day. Furthermore, since our *de novo* sequencing approach does not depend on protease specificity, protease digests from the same sample can potentially be pooled after quenching. Such a multiplexing strategy would reduce both instrument time and analytical costs.

To identify peptides in the InstaNexus workflow, we utilise a leading *de novo* sequencing model and develop a novel assembly workflow that integrates two complementary assembly strategies (greedy and DBG) to extend contig length and improve coverage. We evaluated the identifications from the *de novo* peptide sequencing results and established a methodology where true hits reliably outnumbered false positives while retaining near-complete coverage of target protein sequences. Although both assembly methods share certain parameters, such as identity thresholds and minimum length filters, the greedy strategy offers greater flexibility in handling overlaps, particularly beneficial in noisy or low-coverage datasets. However, this flexibility increases its dependence on parameter tuning. Our assembly pipeline contained few complementary parameters to optimise that might be uniquely different based on the several protein classes that we tested. Our grid search optimization strategy allowed us to optimize the parameters for assembly methods, presenting an optimized scaffold workflow that can be applied to unknown sequences of the same modality. The scoring method we developed considers multiple samples and emphasises additional metrics about sequence assembly rather than relying solely on coverage, integrating coverage with the quality and number of contigs instead. To support broader applicability, we defined a set of optimal parameters for contig and scaffold assembly. These parameters showed minimal variation across different sample types, allowing us to propose a generalizable, consensus configuration (summarized in Supplementary tables 4 and 5). Researchers can use these default settings when applying the workflow to new or unknown samples, particularly in cases where reference-based optimization is not feasible.

These design features distinguish our workflow from existing tools, which offer valuable but more constrained strategies for peptide assembly. MuCS^52^ and ALPS^4^ both employ de Bruijn graph-based approaches, but MuCS lacks parameter optimization and false-positive control, while ALPS depends on homology-based inputs, limiting purely de novo applicability. Stitch^30^ uses template-guided assembly suited to antibody profiling but relies on homologous references and is not applicable to other protein classes. In contrast, our workflow is reference-free, parameter-optimized, and broadly generalizable across protein types and sample contexts.

In the MS analysis of this work, we only utilized HCD fragmentation, which could be supplemented with ETD or other methods to produce complementary peptides that could potentially increase assembly coverage and consensus sequence confidence^37^. Another approach that could be considered would be to use offline fractionation in individual or digest pools to increase detection sensitivity. Both of these approaches, however, would increase sample preparation complexity and analytical costs. Another shortcoming of the methodology used in this work is that, due to their isobaricity, our workflow is unable to discern between isoleucine and leucine, which are therefore treated as interchangeable when mapping assembled contigs to reference scaffolds. While this can increase the number of candidate sequences that need to be validated experimentally, manual inspection of scans can provide additional information about the identity of the residue in a given position via the detection of side chain fragment ions^31^.

In addition to the above limitations, it must be noted that although our workflow consistently produces long contigs for all tested samples, achieving a complete protein sequence with absolute confidence still remains elusive. However, the high coverage and contiguity of target sequences produced by our approach demonstrate the potential to provide truly actionable biological information. Although making a single complete and confident assembly of an antibody is currently lacking, comparative analysis of the generated consensus sequences with known reference sequences yielded 100% accuracy in the CDRs of mAbs and Nbs. Challenges also remain with complex samples, such as oligoclonal and polyclonal protein mixtures, where multiple proteins are present. Nevertheless, our method achieved promising results even in these cases, with performance comparable to that observed for individually assembled mAbs. This challenge was further amplified by the high sequence similarity among mAbs within the mixture, making accurate differentiation inherently more difficult. Other contributing factors to reduced performance with mAbs could be their multi-chain nature and the potential for extensive glycosylation in the heavy chain, a post-translational modification that could be incorporated by future *de novo* peptide sequencing models to improve assembly performance. Additionally, insufficient separation of the mAb chains during sample preparation might also contribute to reduced digestion efficiency, further impacting performance.

Despite the above mentioned challenges, the successful application of our methodology to mAbs represents a significant step toward broader use cases, such as rapid antibody discovery and immune repertoire profiling. For *de novo* designed proteins, direct sequencing enables verification of design fidelity and confirmation of expression. By subjecting binder or antibody pools to selective pressure or target screening and applying *de novo* sequencing, researchers would potentially identify and characterise functional binders with high throughput and quantitative accuracy similar to display technologies^17^, but without the need for prior tagging, library construction, or other molecular biology techniques. This opens the door to direct selection, screening, and sequencing of large antibody and binder libraries at the protein level.

Looking ahead, the next frontier in direct protein sequencing will likely involve its application to increasingly complex biological samples. While this will likely require improving the performance of identification and addition of steps in preparation and data acquisition of our pipeline, the fundamental framework presented here remains broadly applicable and generalizable. Future work could focus on extending this methodology to complex samples and biological systems, enabling sequencing of polyclonal antibody mixtures. It would also unlock applications in metaproteomics, enabling sequencing of complex cellular communities, or sequencing novel enzymes for biotechnological applications. Improvements in *de novo* sequencing models that offer increased accuracy and widely expanded support for diverse post-translational modifications may further enhance our ability to directly sequence proteins by improving alignment accuracy and coverage^40,53^. Together with more comprehensive sample preparation and MS technologies, complete and high-fidelity protein sequencing methodologies may emerge, which can possibly be used for the sequencing of complete proteomes without the need for reference databases.

## Data availability

The InstaNovo checkpoint has been trained on the ProteomeTools datasets, Parts I-III and can be found in the ProteomeXchange Consortium via the PRIDE partner repository with identifiers PXD004732 (Part I), PXD010595 (Part II) and PXD021013 (Part III). The proteomics datasets and database search results generated in this study for BSA, Nbs, mAbs and mBds have been deposited in PRIDE^54^ under dataset identifier PXD066160. Reviewers can access the dataset with username: and password: TeperKWxjeqp. Supplementary data supporting the data preprocessing and analysis performed can be found on Zenodo with doi: 10.5281/zenodo.16417501. The structure of miBd NY1-B04 was resolved and is publicly available in the Protein Data Bank^55^ (PDB-ID: 9NNF).

## Code availability

Inference code and model checkpoints for the base InstaNovo model are available at the InstaNovo GitHub repository https://github.com/instadeepai/instanovo. The InstaNexus pipeline and code can be found in the GitHub repository https://github.com/Multiomics-Analytics-Group/InstaNexus. Custom scripts used for data analysis and visualisation are available upon request and will also be uploaded to a public repository upon publication.

## Acknowledgments

M.R. has been supported by the grant number NNF20CC0035580. K.K. is supported by a Novo Nordisk Foundation Young Investigator Award (grant no. NNF16OC0020670) and a postdoctoral fellowship grant from the Independent Research Fund Denmark (grant no. 4257-00010B). K.K. and M.V.L. also acknowledge support from PRO-MS: Danish National Mass Spectrometry Platform for Functional Proteomics (grant no. 5072-00007B). A.H.L. is supported by a grant from the European Research Council (ERC) under the European Union’s Horizon 2020 research and innovation program (850974), a grant from the Villum Foundation (00025302), and a grant from Wellcome (221702/Z/20/Z). We are grateful to the DTU Bioengineering Proteomics Core Facility for maintenance and running of instruments. Similarly, we thank the Informatics Platform at DTU Biosustain for their support during the optimization of InstaNexus.

## Author contributions

T.P.J., A.S. and K.K. conceived the project. D.S.W., A.L., A.H.L. and T.P.J. provided samples for benchmarking and evaluation. M.W.N., E.L. and M.V.L. performed the enzymatic panel optimization and sample preparation for mass spectrometry. M.W.N. and M.V.L. performed mass spectrometry and searched the data with database search. J.V.G. and K.K. analyzed the mass spectrometry data with de novo peptide sequencing. M.R., A.S. and K.K. developed the software for de novo protein assembly and workflow, with feedback from A.L. and T.P.J. M.R. and P.D.C. performed parameter optimization and tuning for protein assembly. M.R., M.W.N., D.S.W., T.P.J., M.V.L, A.S. and K.K. wrote the manuscript with feedback from A.L., A.H.L., E.M.S and J.V.G. All authors reviewed and approved the final manuscript.

## Additional information

### Competing interests

J.V.G. is an employee of InstaDeep, 5 Merchant Square, London, UK. The other authors declare no competing interests.

### Accession codes

**Accession Table 1.**
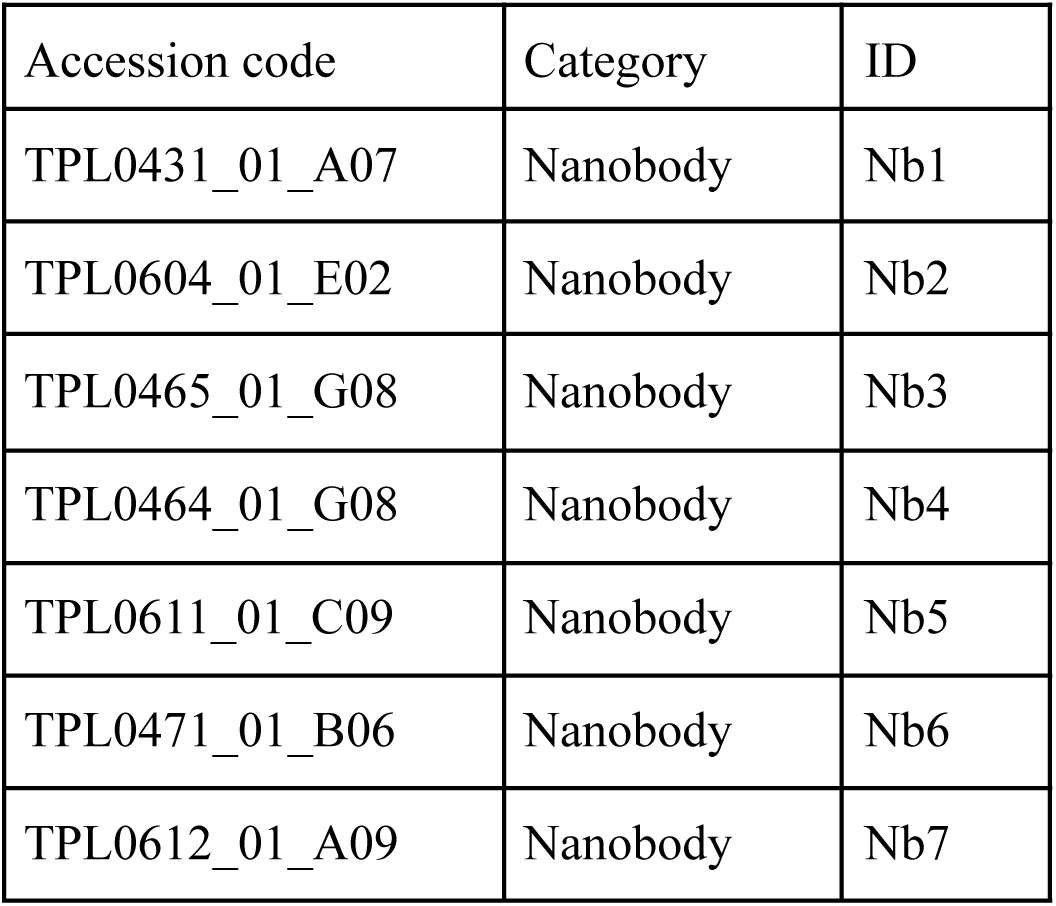
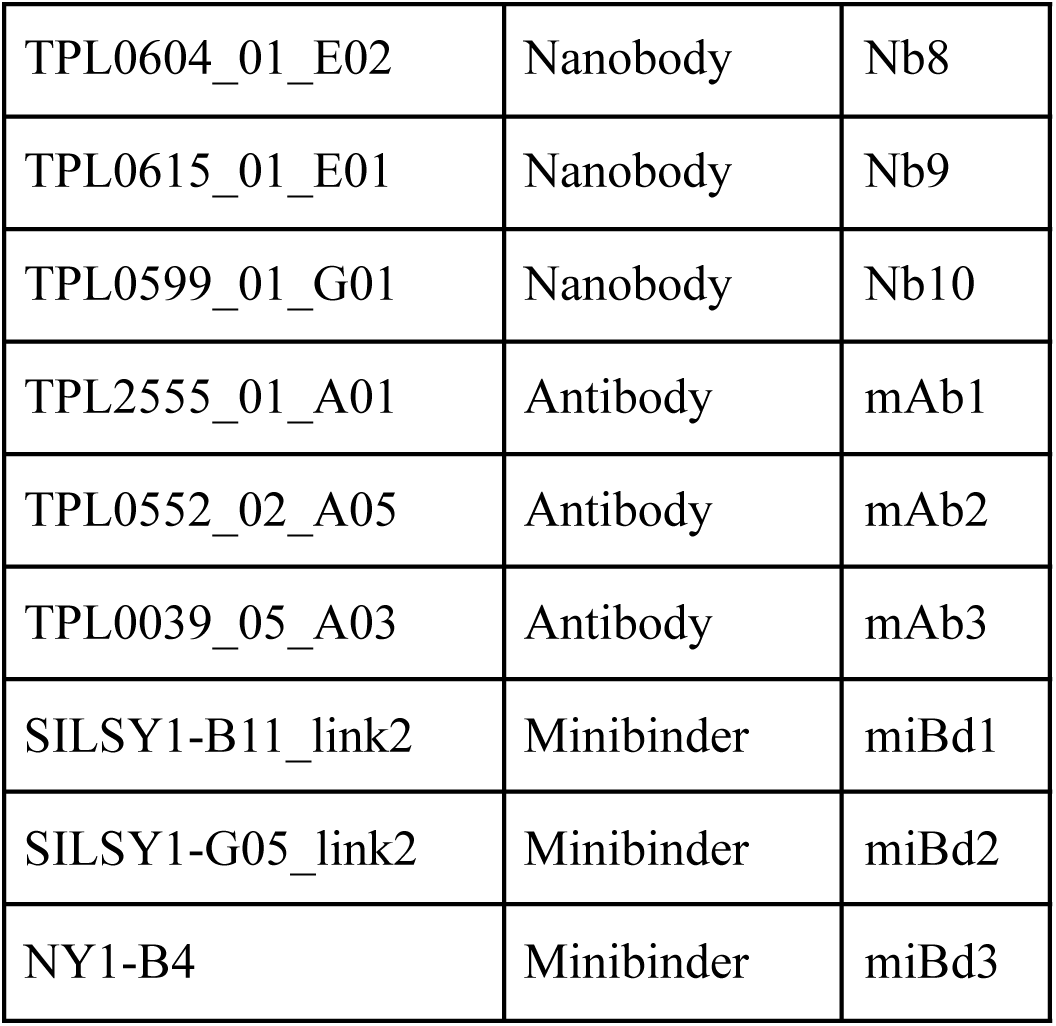
Accession codes for nanobodies, antibodies and minibinders with their IDs in the fasta files used.

## Supplementary information

**Supplementary Figure 1.**
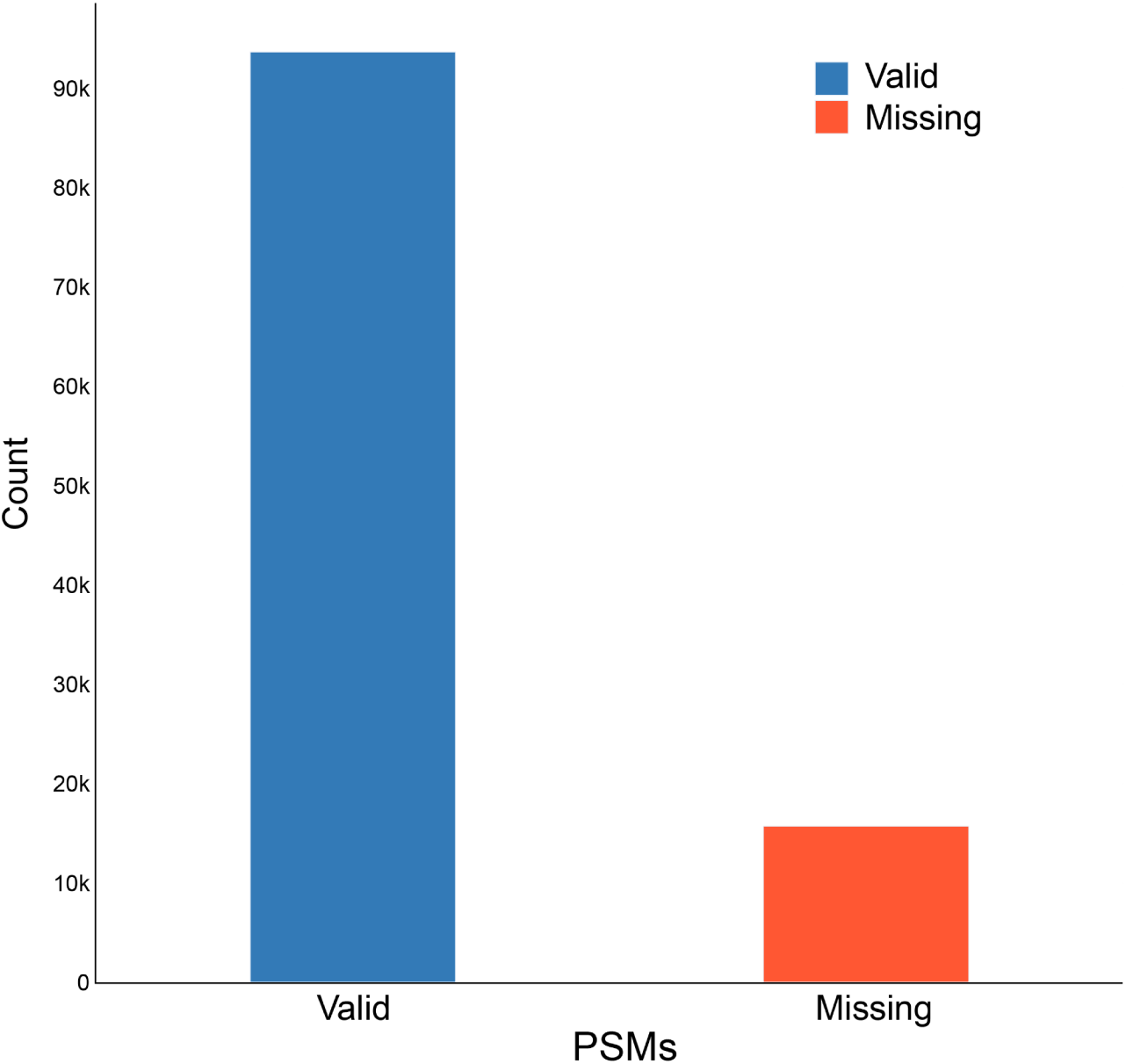
Valid and missing PSMs in BSA. The barplot shows the distribution of peptide-spectrum matches (PSMs) identified during mass spectrometry analysis of BSA. The majority of PSMs are classified as valid (over 90,000), while a smaller fraction are labeled as missing (∼15,000), indicating spectra that were not assigned any confidence score and therefore could not be classified as valid identifications.

**Supplementary Figure 2.**
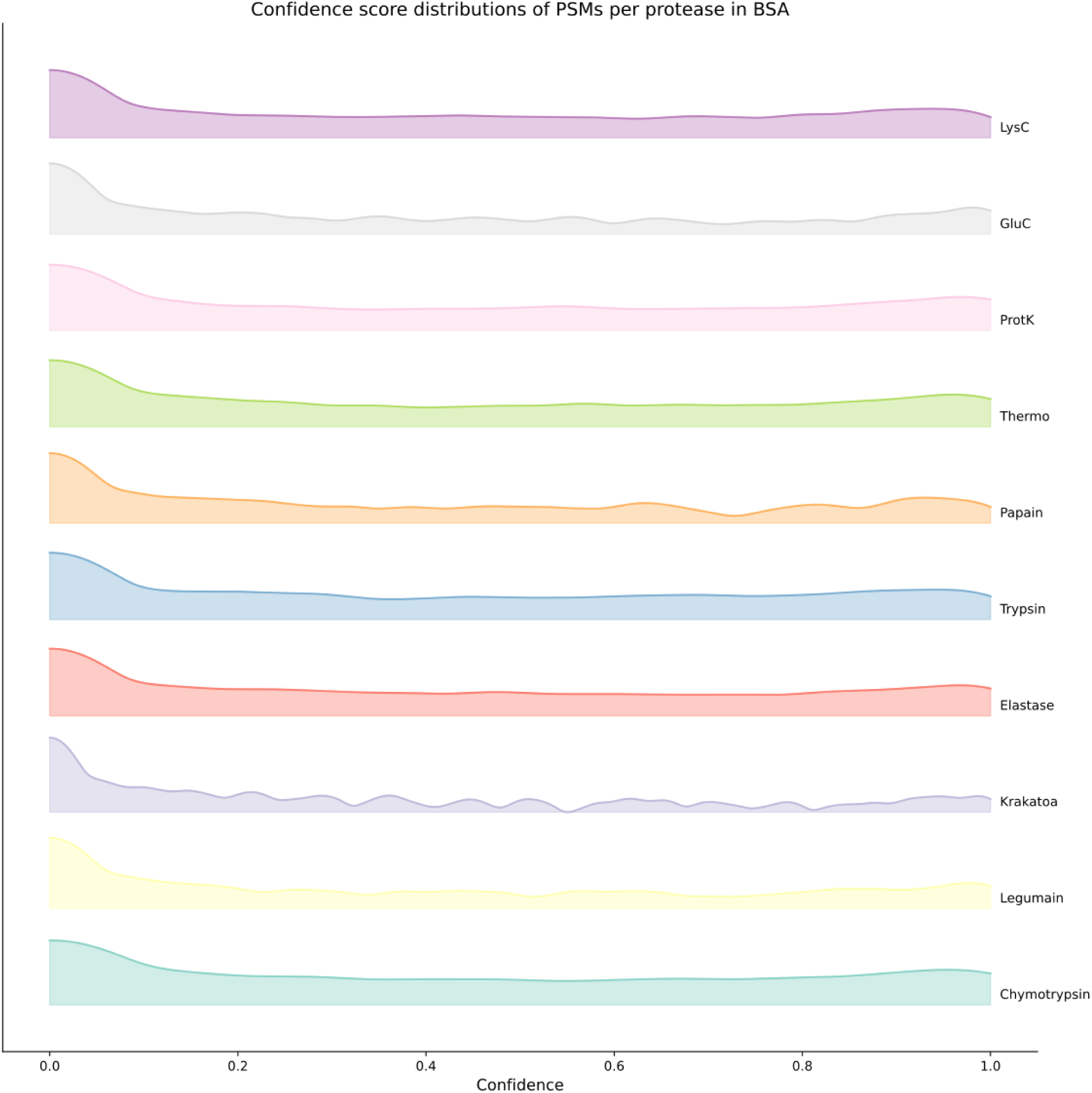
Confidence score distributions across proteases in BSA. The plot shows the kernel density estimates of peptide confidence scores for ten proteases, displayed on a logarithmic (base 10) scale. Each curve represents the distribution of predicted peptide confidence for one protease. Distributions are vertically offset and scaled to allow direct comparison across proteases.

**Supplementary Figure 3.**
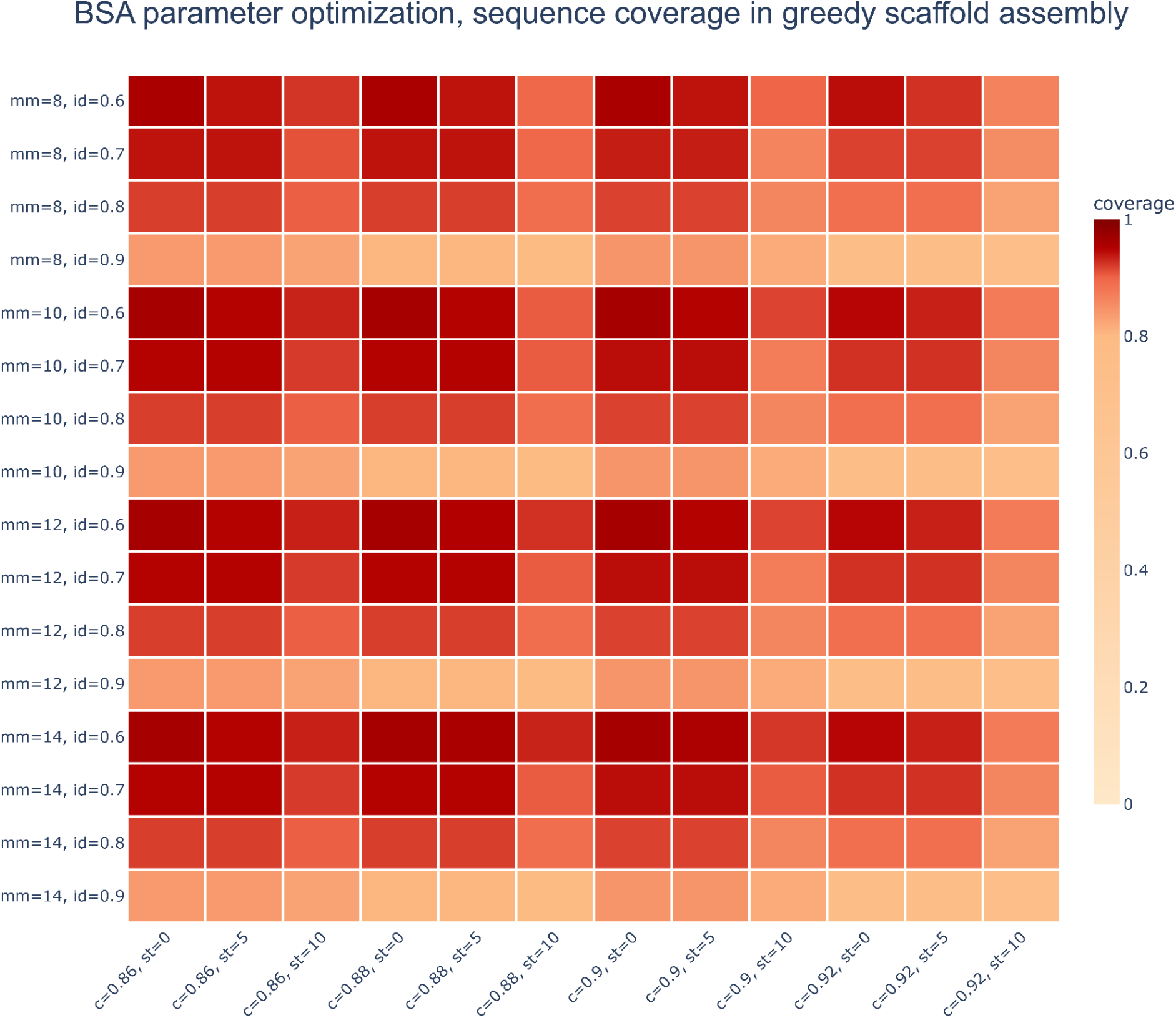
Heatmap of grid search on BSA for greedy scaffold assembly. Heatmap shows the effect of varying assembly parameters on sequence coverage in greedy assembly of BSA. Each row represents a combination of maximum mismatches (mm) and identity threshold (id), while columns represent confidence thresholds (c) and size threshold (st). Coverage values are color-coded, with darker red indicating higher sequence coverage. Higher coverage is observed at intermediate identity thresholds (id = 0.7–0.8) and moderate mismatch tolerance (mm = 12). In contrast, stricter identity thresholds (id = 0.9) and excessive mismatch constraints (mm = 8) result in reduced coverage. Additionally, lower size thresholds (st = 0-5) generally allow for better coverage, whereas higher thresholds (st = 10) tend to reduce coverage by excluding shorter but potentially informative contigs. Confidence threshold (c) values between 0.88 and 0.9 provide consistently high coverage.

**Supplementary Figure 4.**
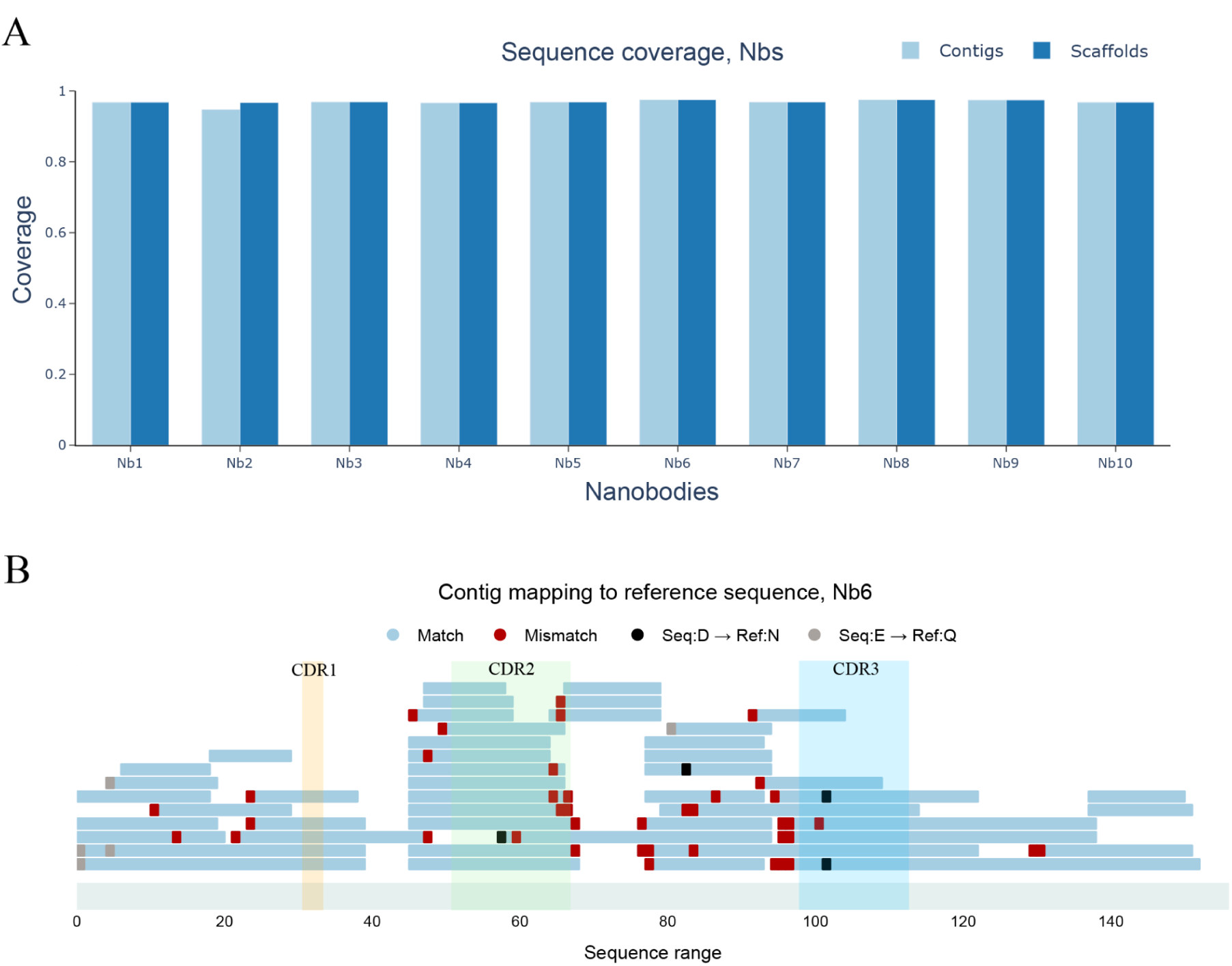
Coverage comparison in nanobody assemblies and contig mapping to a representative nanobody. A) Barplot shows sequence coverage for ten nanobody assemblies (Nb1–Nb10), comparing contig (light blue) and scaffold (dark blue) coverage. All nanobodies display high coverage across both contigs and scaffolds. (B) Mapping of contigs to the reference sequence for nanobody 6. Blue bars represent aligned contigs, with red and black markers indicating mismatches and substitutions. Most of the mismatches are located in the middle of the protein.

**Supplementary Figure 5.**
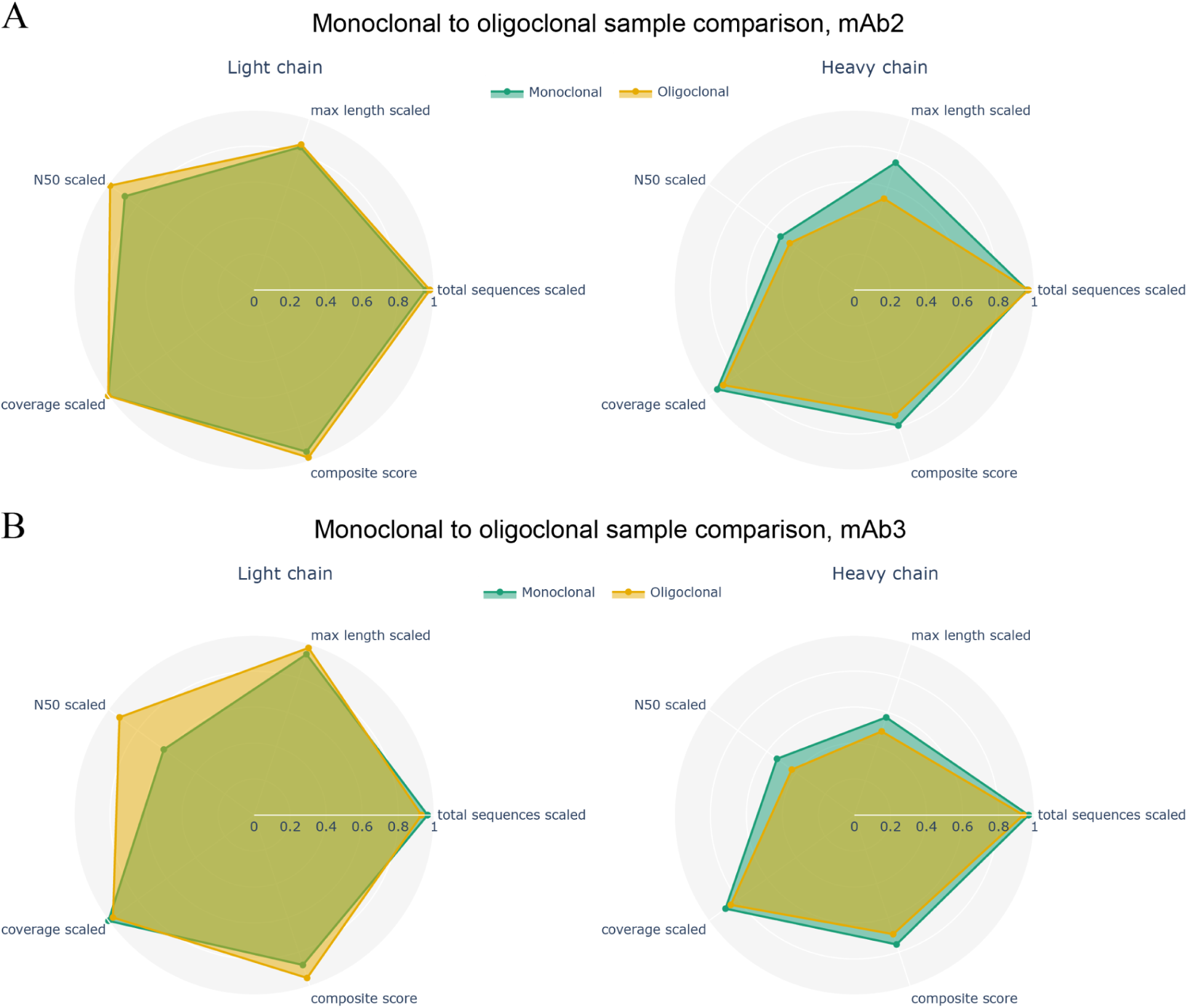
Comparison of monoclonal and oligoclonal antibody sequencing metrics for monoclonal antibody 2 and 3, light and heavy chain. (A) Radar plots showing scaled sequencing metrics for light chain (left) and heavy chain (right) antibody repertoires in antibody 2. Metrics compared include total sequences, maximum length, N50, coverage and a composite score. Both monoclonal (green) and oligoclonal (orange) repertoires show similar profiles across most metrics, with slightly higher heavy chain max length and N50 values in the monoclonal case. (B) Similar comparison for antibody 3. While the light chain metrics are again comparable between conditions, oligoclonal repertoires show increased N50 values compared to monoclonal. Heavy chain metrics remain largely consistent between the two samples.

## Supplementary tables

**Supplementary Table 1.** Assembly metrics for nanobodies using greedy assembly on scaffolds. Summary of sequence assembly statistics for ten nanobody samples (Nb1–Nb10) processed with the greedy assembly method. Nb8 achieved the highest composite score (0.700), associated with high maximum length, coverage, and N50.

| category | run | ass_<br>method | source | total_<br>sequences | total_<br>sequences_<br>scaled | max_<br>length | max_<br>length_<br>scaled | N<br>50 | N50_<br>scaled | cover<br>age | coverage_<br>scaled | composite_<br>score |
| --- | --- | --- | --- | --- | --- | --- | --- | --- | --- | --- | --- | --- |
| nanobodies | Nb1 | greedy | comb_greedy_c0.<br>9_ts10_mo4_mi0<br>.9_mm10 | 35 | 0,868 | 74 | 0,336 | 30 | 0,146 | 0,968 | 0,956 | 0,642 |
| nanobodies | Nb2 | greedy | comb_greedy_c0.<br>86_ts10_mo4_mi<br>0.9_mm14 | 27 | 0,9 | 97 | 0,49 | 38 | 0,208 | 0,947 | 0,929 | 0,666 |
| nanobodies | Nb3 | greedy | comb_greedy_c0.<br>9_ts0_mo3_mi0.<br>9_mm10 | 22 | 0,92 | 90 | 0,443 | 43 | 0,246 | 0,957 | 0,941 | 0,681 |
| nanobodies | Nb4 | greedy | comb_greedy_c0.<br>86_ts10_mo4_mi<br>0.9_mm12 | 18 | 0,936 | 77 | 0,356 | 46 | 0,269 | 0,966 | 0,954 | 0,687 |
| nanobodies | Nb5 | greedy | comb_greedy_c0.<br>88_ts10_mo3_mi<br>0.7_mm14 | 43 | 0,836 | 64 | 0,268 | 37 | 0,2 | 0,968 | 0,957 | 0,649 |
| nanobodies | Nb6 | greedy | comb_greedy_c0.<br>88_ts0_mo4_mi0<br>.9_mm10 | 39 | 0,852 | 94 | 0,47 | 30 | 0,146 | 0,975 | 0,965 | 0,659 |
| nanobodies | Nb7 | greedy | comb_greedy_c0.<br>88_ts10_mo3_mi<br>0.9_mm8 | 33 | 0,876 | 87 | 0,423 | 43 | 0,246 | 0,968 | 0,957 | 0,682 |
| nanobodies | Nb8 | greedy | comb_greedy_c0.<br>88_ts10_mo3_mi<br>0.9_mm14 | 24 | 0,912 | 126 | 0,685 | 36 | 0,192 | 0,975 | 0,966 | 0,7 |
| nanobodies | Nb9 | greedy | comb_greedy_c0.<br>88_ts0_mo3_mi0<br>.9_mm10 | 28 | 0,896 | 93 | 0,463 | 30 | 0,146 | 0,968 | 0,956 | 0,658 |
| nanobodies | Nb10 | greedy | comb_greedy_c0.<br>9_ts10_mo3_mi0<br>.8_mm14 | 39 | 0,852 | 113 | 0,597 | 43 | 0,246 | 0,968 | 0,956 | 0,697 |

**Supplementary Table 2.**
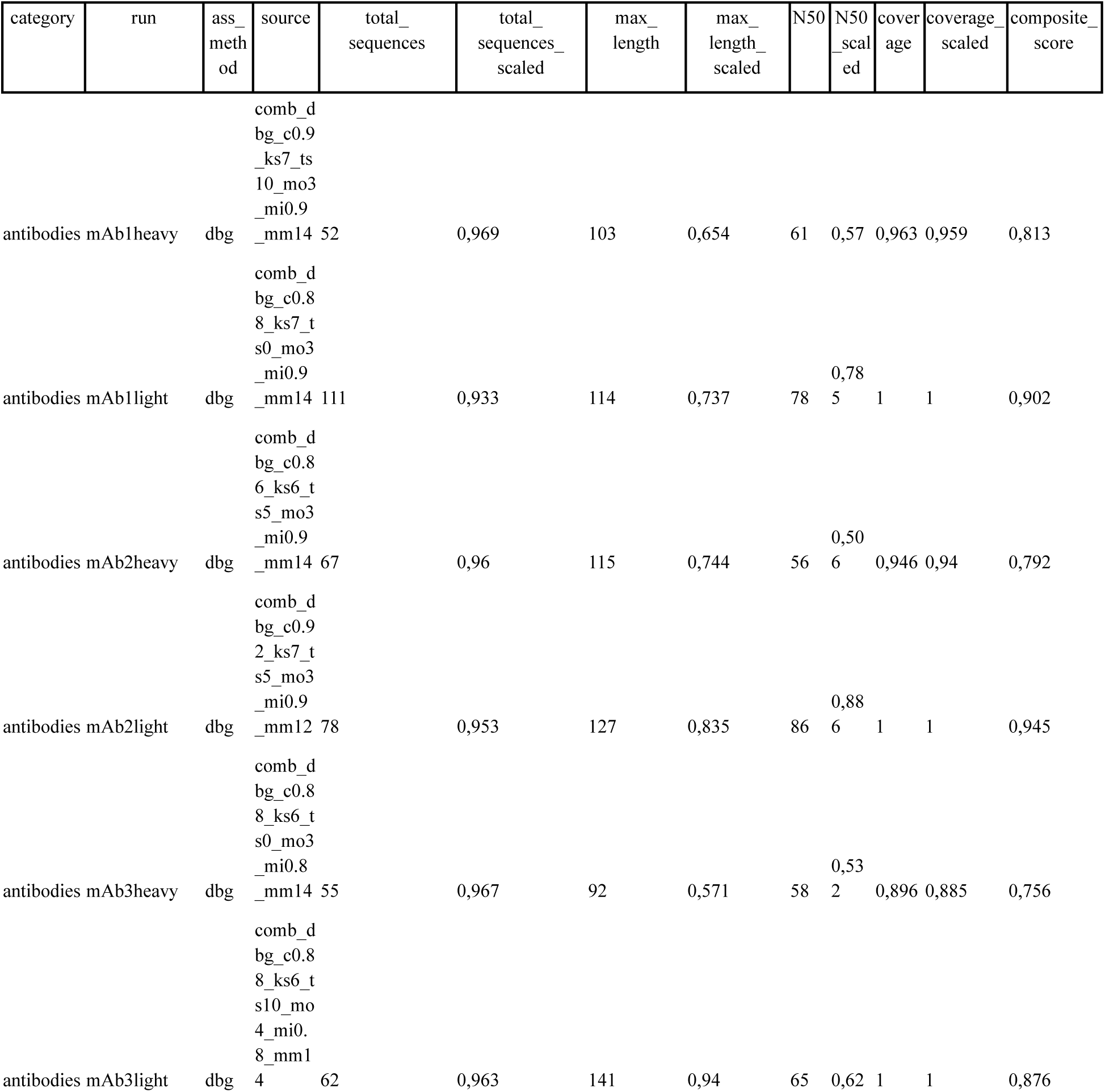
Assembly metrics for mAbs using DBG assembly on scaffolds. Summary of sequence assembly statistics for three mAbs (mAb1, mAb2 and mAb3), each with light and heavy chains assembled using the DBG method. The best-performing light chain was from antibody mAb 2, with the highest composite score (0.945), supported by a long maximum contig (127), high N50 (86), and full coverage (1.0). For heavy chains, the top result was from mAb 1, with a composite score of 0.813.

**Supplementary Table 3.** Assembly metrics for miBds using greedy assembly on scaffolds. Summary of sequence assembly statistics for three miBds, assembled using the greedy method. The best-performing binder was miBd3, with the highest composite score (0.631), associated with a relatively long scaffold (73), N50 (26) and high coverage (0.958).

| category | run | ass_<br>method | source | total_<br>sequences | total_<br>sequences_<br>scaled | max_<br>length | max_<br>length_<br>scaled | N50 | N50_<br>scale<br>d | cover<br>age | coverage_<br>scaled | composite_<br>score |
| --- | --- | --- | --- | --- | --- | --- | --- | --- | --- | --- | --- | --- |
| binders | miBd1 | greedy | comb_greedy_c0.<br>9_ts5_mo3_mi0.9<br>_mm10 | 9 | 0,972 | 53 | 0,195 | 34 | 0,177 | 0,806 | 0,737 | 0,538 |
| binders | miBd2 | greedy | comb_greedy_c0.<br>9_ts0_mo3_mi0.6<br>_mm14 | 33 | 0,876 | 37 | 0,087 | 21 | 0,077 | 0,798 | 0,725 | 0,482 |
| binders | miBd3 | greedy | comb_greedy_c0.<br>92_ts10_mo3_mi<br>0.6_mm14 | 21 | 0,924 | 73 | 0,329 | 26 | 0,115 | 0,958 | 0,943 | 0,631 |

**Supplementary Table 4.** Mode values of DBG assembly parameters observed across different protein categories. Mode values of key parameters used during greedy scaffold-based assembly for four protein types: BSA, Nbs, miBds, and mAbs. Most parameters show high consistency across categories, with identical mode values observed for minimum identity (0.9 in three out of four categories), maximum mismatches (14 in all but BSA), k-mer size (6 in three categories), and minimum overlap (3 across all categories).

| Category | Confidence | Size threshold | Minimum identity | Maximum mismatches | K-mer size | Minimum overlap |
| --- | --- | --- | --- | --- | --- | --- |
| BSA | 0.86 | 0 | 0.9 | 12 | 6 | 3 |
| Nanobodies | 0.9 | 5 | 0.9 | 14 | 6 | 3 |
| Minibinders | 0.88 | 0 | 0.6 | 14 | 7 | 3 |
| Antibodies | 0.88 | 0 | 0.9 | 14 | 6 | 3 |

**Supplementary Table 5.** Mode values of greedy assembly parameters observed across different protein categories. Mode values of key parameters used during greedy scaffold-based assembly for four protein types: BSA, Nbs, miBds, and mAbs. Most parameters are consistent across categories, with minimum overlap fixed at 3 in all cases, and identical confidence (0.88) observed for three out of four categories. Size threshold and minimum identity also show conserved values in most categories, with slight variation for miBds.

| Category | Confidence | Size threshold | Minimum identity | Max mismatches | Minimum overlap |
| --- | --- | --- | --- | --- | --- |
| BSA | 0.88 | 10 | 0.6 | 14 | 3 |
| Nanobodies | 0.88 | 10 | 0.9 | 10 | 3 |
| Minibinders | 0.9 | 0 | 0.6 | 14 | 3 |
| Antibodies | 0.88 | 10 | 0.9 | 10 | 3 |

## Notes

https://zenodo.org/records/16417502

https://github.com/Multiomics-Analytics-Group/InstaNexus

